# Dysregulation of amino acid metabolism upon rapid depletion of cap-binding protein eIF4E

**DOI:** 10.1101/2023.05.11.540079

**Authors:** Paige D. Diamond, Nicholas J. McGlincy, Nicholas T. Ingolia

**Affiliations:** Department of Molecular and Cell Biology, University of California, Berkeley; Center for Computational Biology and California Institute for Quantitative Biosciences, University of California, Berkeley

## Abstract

Protein synthesis is a crucial but metabolically costly biological process that must be tightly coordinated with cellular needs and nutrient availability. In response to environmental stress, translation initiation is modulated to control protein output while meeting new demands. The cap-binding protein eIF4E—the earliest contact between mRNAs and the translation machinery—serves as one point of control, but its contributions to mRNA-specific translation regulation remain poorly understood. To survey eIF4E-dependent translational control, we acutely depleted eIF4E and determined how this impacts protein synthesis. Despite its essentiality, eIF4E depletion had surprisingly modest effects on cell growth and protein synthesis. Analysis of transcript-level changes revealed that long-lived transcripts were downregulated, likely reflecting accelerated turnover. Paradoxically, eIF4E depletion led to simultaneous upregulation of genes involved in catabolism of aromatic amino acids, which arose as secondary effects of reduced protein biosynthesis on amino acid pools, and genes involved in the biosynthesis of amino acids. These futile cycles of amino acid synthesis and degradation were driven, in part, by translational activation of *GCN4*, a transcription factor typically induced by amino acid starvation. Furthermore, we identified a novel regulatory mechanism governing translation of *PCL5,* a negative regulator of Gcn4, that provides a consistent protein-to-mRNA ratio under varied translation environments. This translational control was partial dependent on a uniquely long poly-(A) tract in the *PCL5* 5’ UTR and on poly-(A) binding protein. Collectively, these results highlight how eIF4E connects translation to amino acid homeostasis and stress responses and uncovers new mechanisms underlying how cells tightly control protein synthesis during environmental challenges.

## Introduction

Protein synthesis, a vital step in gene expression and a significant biosynthetic process, is tightly regulated in response to nutrient status and environmental cues (Crawford and Pavitt, 2019; Hershey et al., 2012; Liu and Qian, 2014; Sonenberg and Hinnebusch, 2009). Often, this regulation affects translation initiation, a rate-limiting step and a point of commitment to protein synthesis (Shah et al., 2013). In eukaryotes, deeply conserved pathways modulate overall protein synthesis levels by controlling recognition of the distinctive 5’ cap on mRNAs and recruitment of the initiator tRNA to begin translation. These pathways also control transcript-specific expression, and sequence features in mRNAs play important roles in determining levels of protein synthesis (Park et al., 2011; Sen et al., 2016; Thoreen et al., 2012; Zinshteyn et al., 2017). Inhibitory upstream open reading frames (uORFs) divert initiation machinery and ribosomes from the main ORF, thus inhibiting translation in a context-dependent manner that can be relieved in stress (Brar et al., 2012; Calkhoven et al., 2000; Lu et al., 2004; Young and Wek, 2016). Translation of *GCN4,* a master regulator of amino acid starvation response in yeast, is a prime example of such uORF-dependent translation (Hinnebusch, 2005; Mueller and Hinnebusch, 1986). In response to amino acid deprivation, the kinase Gcn2 phosphorylates the translation initiation factor eIF2α (Dever et al., 1992; Hinnebusch, 2005), inhibiting bulk translation by decreasing the availability of eIF2 complex with initiator tRNA. *GCN4*, along with other stress-responsive proteins, overcome this inhibition to ensure the production needed to respond to stress. However, the mechanisms by which these transcripts are differentially translated is not fully understood.

The cap-binding protein, eIF4E, makes the first contact between a transcript and the translation initiation machinery during a typical initiation event. By recognizing the 5’m^7^G cap, a hallmark feature of eukaryotic mRNAs, eIF4E plays a central role in the canonical, cap-dependent pathway for translation initiation (Marcotrigiano et al., 1997; Matsuo et al., 1997; von der Haar et al., 2004). It also regulates mRNA stability by preventing decapping and degradation, perhaps through steric hindrance of decapping machinery (Vilela, 2000). Differential binding of cap-binding proteins to specific mRNAs regulates translational responses to environmental stresses in mammalian systems. Transcripts with 5’-terminal oligopyrimidine (5’TOP) tracts, which encode ribosomal proteins and elongation factors, are particularly sensitive to growth conditions and amino acid availability (Fonseca et al., 2015; Meyuhas and Kahan, 2015). In starvation, La-related protein 1 (LARP1) binds the caps of 5’TOP mRNAs to impede access of eIF4E to these specific transcripts (Lahr et al., 2017; Philippe et al., 2020). While *Saccharomyces cerevisiae* lacks this mode of regulation, yeast eIF4E is encoded by *CDC33* and was first identified through its cell cycle arrest phenotype (Reid and Hartwell, 1977). The link between eIF4E and cell-cycle progression has been attributed to translational control of the G1 cyclin 3 (Cln3), which drives the transition from G1 to S phase (Danaie et al., 1999). Furthermore, differential engagement of eIF4E across yeast mRNAs has been reported both *in vitro* by smFRET and *in vivo* by RNA-immunoprecipitation sequencing (Çetin and O’Leary, 2022; Costello et al., 2015). These findings raise the broader question, whether eIF4E might preferentially promote translation of certain transcripts, based features of the mRNAs, to control functionally related genes.

To understand the role of eIF4E in translation regulation, we investigated its mRNA-specific effects by profiling changes in gene expression and cell physiology after rapid depletion of eIF4E protein. Depletion of eIF4E preferentially destabilized long-lived mRNAs and reduced bulk protein synthesis. These conditions strongly induced genes involved in the catabolism of aromatic amino acids as secondary effects of reducing protein biosynthesis and the resultant rebalancing of amino acid pools. Furthermore, reductions in eIF4E levels caused translational activation of *GCN4*, a key regulator of stress responsive translation that is typically induced by amino acid starvation. Interestingly, translation of *GCN4* occurred through a non-canonical mechanism instead of relying on the well-established effects of eIF2α phosphorylation. Additionally, we noted translational repression of *PCL5*, a negative regulator of Gcn4, in eIF4E depleted cells, which may have further contributed to imbalanced amino acid pools. This work provided new insights into the regulation of Gcn4 activity through Pcl5-mediated feedback control and highlight the role of yeast eIF4E in control of nutrient-responsive gene regulatory networks.

## Results

### eIF4E depletion destabilizes short, stable transcripts with few direct effects on translation

To investigate the contribution of the cap-binding protein eIF4E to the specificity of translation initiation across the transcriptome, we acutely depleted eIF4E (encoded by *CDC33* in *S. cerevisiae*) and measured the translational consequences. Prior studies relied on temperature-sensitive alleles, which require shifts to non-permissive temperatures that inhibit translation initiation even in wild-type cells and may have complex and incomplete molecular effects (Groušl et al., 2009). To avoid complications associated with temperature sensitivity, we depleted eIF4E using the auxin-inducible degron (AID) system, which allows conditional depletion of a target protein via degradation by the ubiquitin-proteasome system (Kubota et al., 2013; Nishimura et al., 2009) (**Figure 1A**), Degradation of eIF4E was rapidly induced by adding the auxin indole-3-acetic acid (IAA), and <10% of protein remained after 60 minutes (**Figure 1C, S1G**). To measure changes to bulk protein synthesis levels following eIF4E depletion, we carried out metabolic labeling of nascent peptides through addition of methionine analog, L-homopropargylglycine (HPG) for 120 minutes (Wiltschi et al., 2008) (**Figure 1B, 1A-F**). We then measured nascent peptide labeling by flow cytometry, after a fluorophore was added to incorporated HPG using click chemistry. We observed an approximate 50% reduction in nascent peptide labeling after 1 hour IAA treatment relative to DMSO treated cells (**Figure 1B**). This reduction in bulk protein synthesis was also supported by polysome profiling after 1 hour of IAA treatment (**Figure S1H**), which showed an increased proportion of free ribosomal subunits and monosomes and a depletion of polysomes, indicative of a reduction in translation initiation.

**Figure 1:**
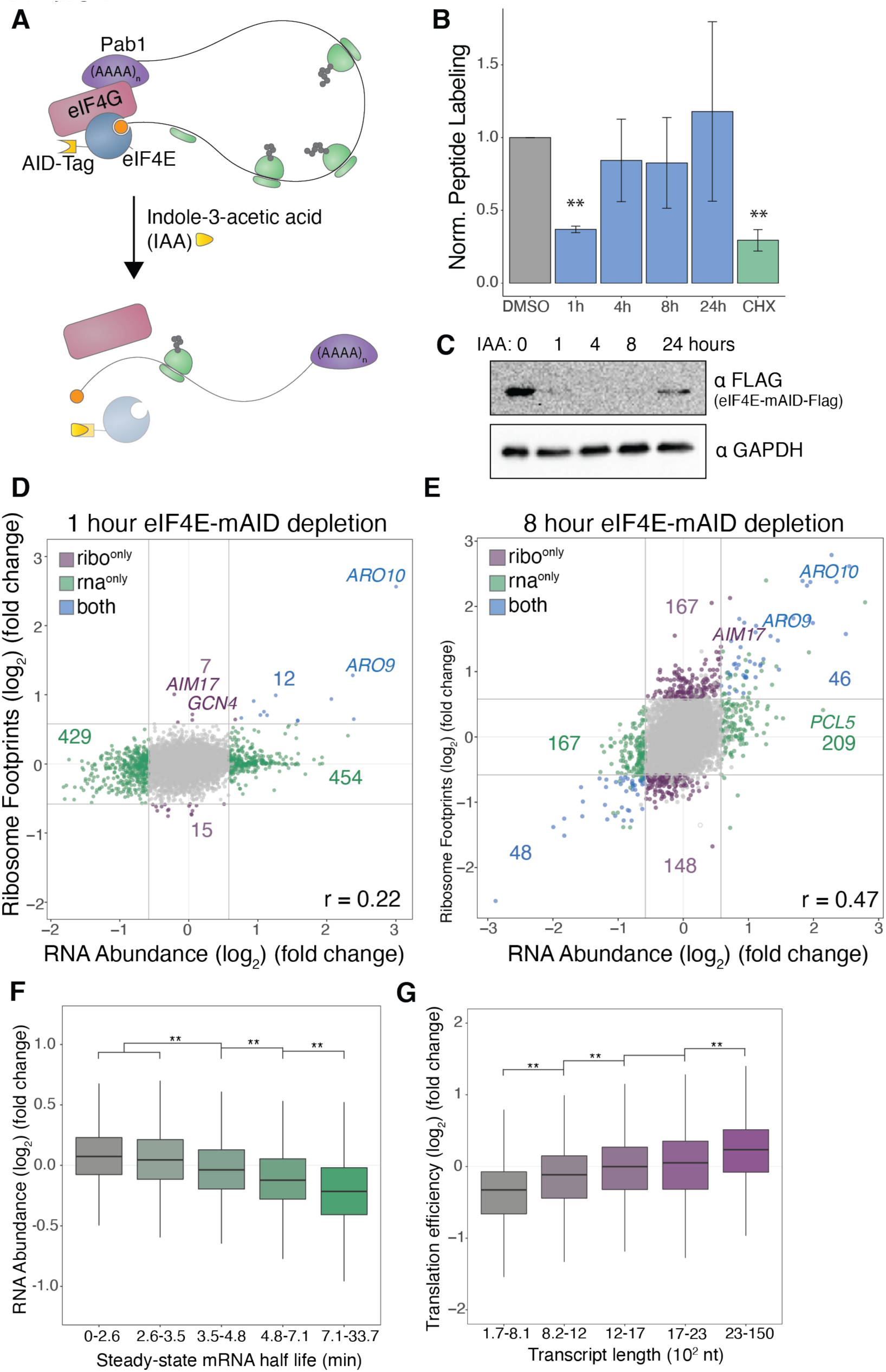
Transcript-level sensitivities to eIF4E depletion. **(A)** Schematic of auxin-inducible degron (AID) tagging and conditional depletion of eIF4E. **(B)** Bulk translation measured by nascent peptide metabolic labeling in eIF4E depleted cells. Indole-3-acetic acid (IAA) or cycloheximide (CHX) was added for indicated durations and maintained during a 2-hour labeling period with L-Homopropargylglycine (HPG). Median intensities of the Alexa Fluor^TM^ 488 (HPG) signal normalized to that of DMSO treated cells. (**) represents p < 0.05 calculated by Student’s t test. Error bars represent standard error of the mean, n=2. **(C)** Western blot for eIF4E-mAID-Flag expression levels over the course of IAA depletion. **(D)** Differential expression after 1 hour of eIF4E depletion measured by RNA-seq and ribosome profiling. IAA-treated cells are compared with DMSO-treated controls. Color represents significant (adjusted p-value < 0.05) and substantial (absolute fold-change (log2) > 0.58) changes. Correlation coefficient (Pearson’s) calculated between log2 fold changes of RPF and RNA abundance. **(E)** Same as (D) except 8-hour IAA treatment. **(F)** Box plots of RNA abundance fold change (log2) for transcripts grouped based on steady-state mRNA half-life^45^. (**) indicates p < 0.05; one-way ANOVA test followed by Tukey’s HSD test. **(G)** Box plots of translation efficiency fold change (log2) for transcripts grouped based on length. (**) indicates p < 0.05; one-way ANOVA test followed by Tukey’s HSD test.

To determine the impact of eIF4E depletion globally on mRNA stability and protein synthesis, we measured changes in mRNA abundance and translation using RNA-sequencing and ribosome profiling (Ingolia et al., 2009). By comparing cells treated with IAA for 1 hour to DMSO control cells, we hoped to capture the most immediate gene expression changes attributed to eIF4E depletion, rather than the accumulation of secondary effects. We observed widespread changes in the transcriptome following eIF4E depletion; 454 transcripts were significantly upregulated and 429 transcripts were downregulated (**Figure 1D, S2A,D**). Surprisingly, these transcript-level changes in RNA expression generally were not reflected in the ribosome occupancy measurements. Lack of strong correlation between these data suggested that transcriptional changes had not propagated through translation. We were interested in measuring gene expression after a longer depletion to see if this discrepancy persisted. After 8 hours of IAA treatment, when eIF4E levels remain undetectably low (**Figure 1C**), we see greater agreement in the RNA abundance and ribosome occupancy measurements (**Figure 1E, S3A-B**). The improved correlation at 8 hours suggested that the disparities we saw after 1 hour of IAA treatment reflected pre-steady-state changes in translation. Although we saw larger changes in ribosome occupancy affecting more genes at the later timepoint, the total level of translation as measured by the HPG metabolic labeling assay recovered after 4 hours of eIF4E depletion despite unmeasurable levels of eIF4E (**Figure 1B,C**).

We hypothesized that the sensitivity of transcripts to eIF4E depletion may be influenced by certain RNA attributes, such as poly(A) length or 5’ UTR structure, based on known interactions between eIF4E, eIF4G, eIF4A, and Pab1 (**Figure S2G, S3G**). Correlations between published mRNA features and the changes we observed following eIF4E depletion yielded a relationship between mRNA stability and sensitivity to eIF4E depletion (**Figure 1F, S2B-C, S3E**) (Chan et al., 2018). Long-lived transcripts (those with a steady-state half-life greater than ∼5 minutes) generally decreased in abundance following eIF4E depletion. The correlation between mRNA half-life and change in abundance following eIF4E depletion was also present after 1-hour IAA treatment (**Figure S2C**). Thus, these stability-related changes happened relatively quickly in response to eIF4E depletion, suggesting that eIF4E preferentially stabilizes those transcripts under unperturbed conditions. We also calculated correlations between RNA attributes and eIF4E-dependent changes in translation efficiency, which is measure calculated from ribosome footprints density changes given underlying RNA expression patterns. Changes in translation efficiency following eIF4E depletion most strongly correlated with transcript length (**Figure 1G, S2E-F, S3F**). Depletion of eIF4E reduces the translation efficiency of short mRNAs and depresses the length bias of translation efficiency. Overall, our findings suggested eIF4E promoted the expression of stable, short transcripts.

### Reduced translation drives aromatic amino acid accumulation and ARO10 induction

While the above analysis revealed overall features associated with sensitivity to eIF4E depletion, the two genes most strongly upregulated following 1-hour eIF4E depletion, both at the level of RNA expression and ribosome occupancy, were *ARO9* and *ARO10*. These genes encode enzymes involved in the catabolism of aromatic amino acids via the Ehrlich pathway, which facilitates the utilization of aromatic amino acids as nitrogen sources under nitrogen-limiting conditions (**Figure 1D**) (Kradolfer *et al*. 1982; Vuralhan *et al*. 2003; Hazelwood *et al*. 2008). To better understand the gene regulatory network controlling *ARO9* and *ARO10* expression and its connection to cap-dependent translation factors, we established a reporter system to measure changes in *ARO10* transcription and translation. In this experimental setup, we fused a synthetic transcription factor (ZEM) to the endogenous *ARO10* CDS, separated by a self-cleaving P2A peptide (Aranda-Díaz et al., 2017). We contemporaneously introduced an RFP reporter under the control of an orthogonal promoter, p(Z), whose transcriptional output is proportional to ZEM protein abundance. Thus, we quantified *ARO10* transcription via *ARO10-ZEM* mRNA abundance, and indirectly reported on translation of *ARO10* by monitoring expression of RFP from the p(Z) promoter driven by the ZEM protein (**Figure S4A**). Our indirect reporter was induced to a similar degree by eIF4E depletion as endogenous *ARO10*, although its response was delayed slightly because reporter induction requires synthesis and nuclear import of the synthetic transcription factor (**Figure 2A,B**). Having established that this system recapitulated *ARO10* induction following eIF4E depletion, we set out to identify the regulatory network underlying this response.

**Figure 2:**
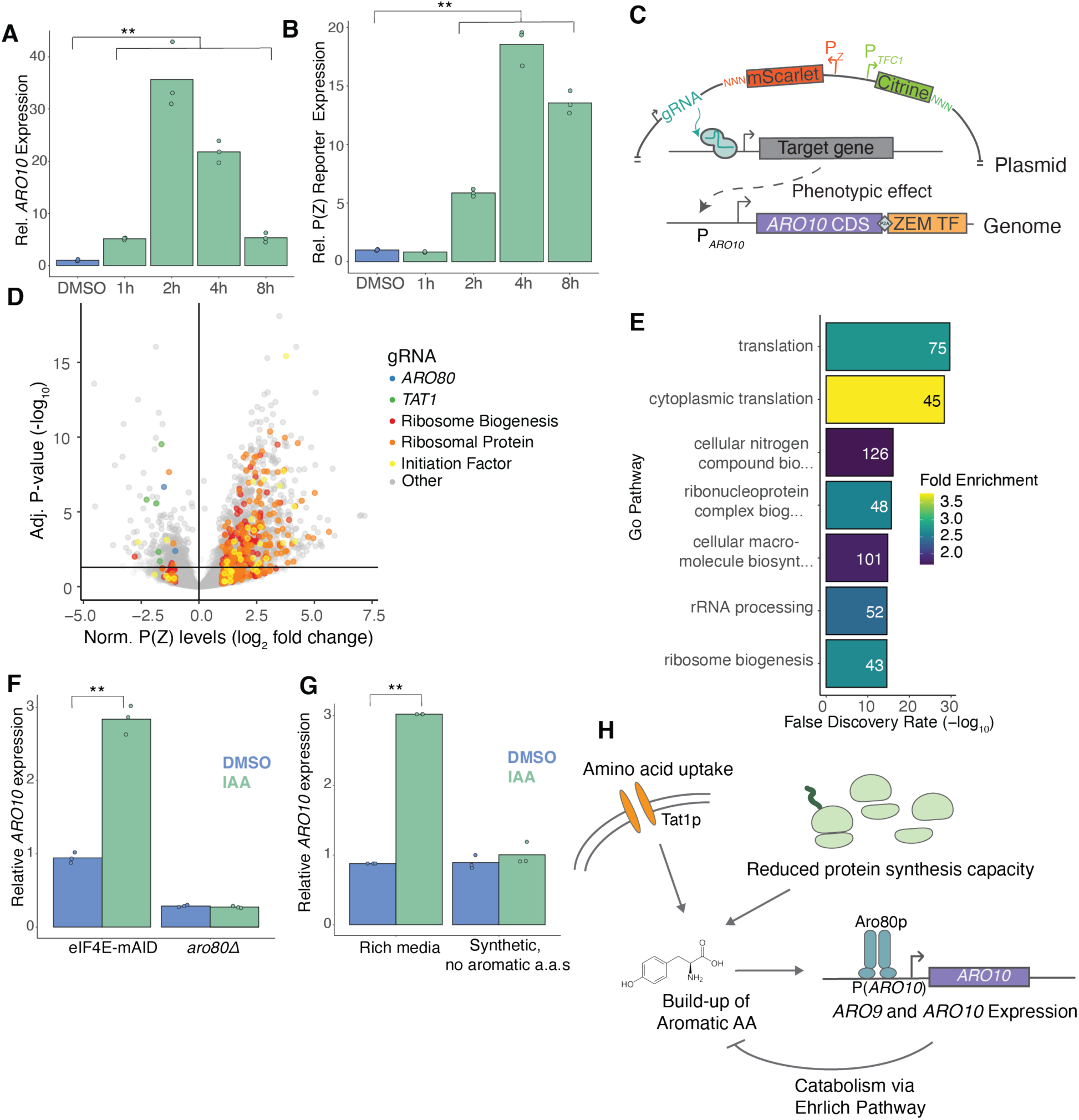
CiBER-seq profile *ARO10* expression regulation. **(A)** RT-qPCR of *ARO10* expression over the course of eIF4E-AID depletion. (**) represents p < 0.05 calculated by Student’s t test. **(B)** RT-qPCR of P(Z) reporter expression over the course of eIF4E-AID depletion. (**) represents p < 0.05 calculated by Student’s t test. **(C)** Schematic of CiBER-seq screen design. **(D)** Genome-wide CiBER-seq results showing fold-change (log2) in P(Z) reporter abundance, relative to P(*TFC1*) reporter levels, for each gRNA. Line indicates significance cutoff (adjusted p-value < 0.05). Color represents gene category or identity. **(E)** GO analysis for genes targeted by gRNAs that up-regulated P(Z) expression. gRNAs were filtered for fold-change (log2) > 1 and adjusted p-value < 0.05. The most statistically significant entries were chosen and narrowed based on percentage of overlapping genes. **(F)** RT-qPCR of *ARO10* expression following 1-hour of eIF4E-AID depletion in *ARO80* and *aro80Δ* cells. (**) represents p < 0.05 calculated by Student’s t test. **(G)** RT-qPCR of ARO10 expression following 1-hour of eIF4E-AID depletion in rich media and synthetic medium without aromatic amino acids. (**) represents p < 0.05 calculated by Student’s t test. **(H)** A model for *ARO10* upregulation and subsequent feedback in response to translational stress and aromatic amino acid availability in the medium.

To uncover the genetic regulatory network driving *ARO10* expression and the connections to eIF4E depletion, we used a genome-wide CRISPRi approach, based on CRISPRi with barcoded expression readout (CiBER-seq), that couples expression of our *ARO10* reporter protein via a unique RNA barcode to a specific genetic perturbation (Muller et al., 2020) (**Figure 2C**). We generated a library of approximately 48,000 gRNAs, each of which was linked to expressed RNA barcodes that could be tracked by next-generation sequencing. We measured barcode abundance from transcripts driven by the P(Z) reporter normalized against a paired barcode expressed from the housekeeping *TFC1* promoter, which allowed us to correct for knockdown effects that affect overall RNA transcription or cellular fitness. We sequenced the RNA barcodes before and after gRNA induction to understand how knockdown of factors changed *ARO10* reporter expression (Ashuach et al., 2019). As expected, guides targeting *ARO10* or *GAL1* (from which P(Z) is derived) were some of the strongest repressors of *ARO10* reporter expression, and we further validated the screen results by individually testing both a gRNA that reduced P(Z)-reporter when induced (*PAB1*) and a gRNA against *TIF34* which increases *ARO10* expression when induced (**Figure S4E,F**).

Our genome-wide screen revealed numerous gRNAs targeting ribosomal protein genes (RPGs) or genes involved in ribosome biogenesis (RiBi) that activated our *ARO10* reporter (**Figure 2D**). This pattern was reflected in GO analysis for gRNAs that significantly increased reporter expression (**Figure 2E**). This suggested to us that *ARO10* induction was not a unique response feature of eIF4E depletion, but instead was upregulated by any reduction of general translation machinery. Conversely, we found gRNAs targeting *ARO80*, a transcription factor known to regulate *ARO10*, that reduced *ARO10* expression (**Figure 2D**) (Lee and Hahn, 2013). Thus, we measured the *ARO10* response to eIF4E depletion in a strain lacking *ARO80* and found that, indeed, a*ro80*Δ abrogated *ARO10* induction (**Figure 2F**).

We also found several gRNAs that significantly reduced *ARO10* expression by targeting components of the SPS-sensing pathway, which regulates the transcription of amino acid permeases in response to extracellular amino acids (Ljungdahl, 2009) (**Figure S4H,I**). While *ARO10* could be a direct, but unreported, transcriptional target of the SPS pathway, we considered whether this could instead reflect an indirect effect. To explore this possibility, we searched for gRNAs against annotated targets of SPS transcription factors Stp1 or Stp2 that significantly reduced *ARO10* reporter expression (Eckert-Boulet et al., 2004). Knockdown of *TAT1*, a known target of Stp1 and Stp2 that encodes an amino acid transporter for tyrosine, leucine, isoleucine and valine, reduced *ARO10* expression (Bajmoczi et al., 1998; Schmidt et al., 1994). Notably, Tat1 imports the major substrates for Aro9 and Aro10, and Aro80 is allosterically activated by tryptophan (Lee and Hahn, 2013). Thus, SPS control of aromatic amino acid import may explain these effects on *ARO10* expression. To test the possibility that the import of extracellular amino acids mediated *ARO10* expression, we grew cells in synthetic media lacking aromatic amino acids and found that this condition indeed mitigated the induction of *ARO10* after eIF4E depletion (**Figure 2G**). Taken together, these findings suggest that the buildup of aromatic amino acids imported by Tat1 but not needed for protein synthesis is sensed by Aro80, which upregulates *ARO9* and *ARO10* to catabolize them (**Figure 2H**).

We reasoned that this catabolism might mitigate toxicity induced by excess aromatic amino acids or produce a metabolite needed for adaptive responses to reductions of protein synthesis. However, neither deletion of *ARO9* nor of *ARO80* caused any difference in growth rate upon eIF4E depletion. (**Figure S5A,B**). Thus, the induction of the Ehrlich pathway enzymes is not required in cells depleted of eIF4E.

Surprisingly, long-term eIF4E depletion led to the up-regulation of *ARO1* (**Figure S3B**), which encodes a multi-functional chorismite biosynthesis enzyme required to produce aromatic amino acids (Duncan et al., 1988). Induction of this biosynthetic enzyme, along with continued expression of the opposing catabolic pathway, suggested that eIF4E depletion led to futile or gratuitous metabolism. Indeed, we found that cells depended on aromatic amino acid biosynthesis after eIF4E depletion, as shown by slowed growth of *aro1Δ* cells (**Figure S5C**). Furthermore, the negative synthetic interaction between eIF4E depletion and *aro1Δ* was partially rescued by supplementing media with additional aromatic amino acids (**Figure S5D**). Combined, these results show that eIF4E depletion disrupts amino acid homeostasis.

### eIF4E depletion activates *GCN4* translation in *GCN2*-independent manner

Indeed, we saw broad up-regulation of amino acid biosynthesis pathways following 8 hours of eIF4E depletion, indicating a breakdown in coordination between amino acid levels and the regulation of genes involved in amino acid biosynthesis (**Figure 3A**). These biosynthetic genes comprise the regulon of Gcn4, a transcription factor that is itself translationally induced in response to amino acid starvation (Hinnebusch, 2005). Indeed, RNA abundance of Gcn4 target genes increased after 8 hours of IAA treatment compared to non-target genes (**Figure 3B**). We also found a modest but significant translational upregulation of Gcn4 (log_2_ fold-change = 0.61, adj. *p*-value = 1.1×10^-4^) after 1 hour of eIF4E depletion, as measured by ribosome occupancy (**Figure 3C**).

**Figure 3:**
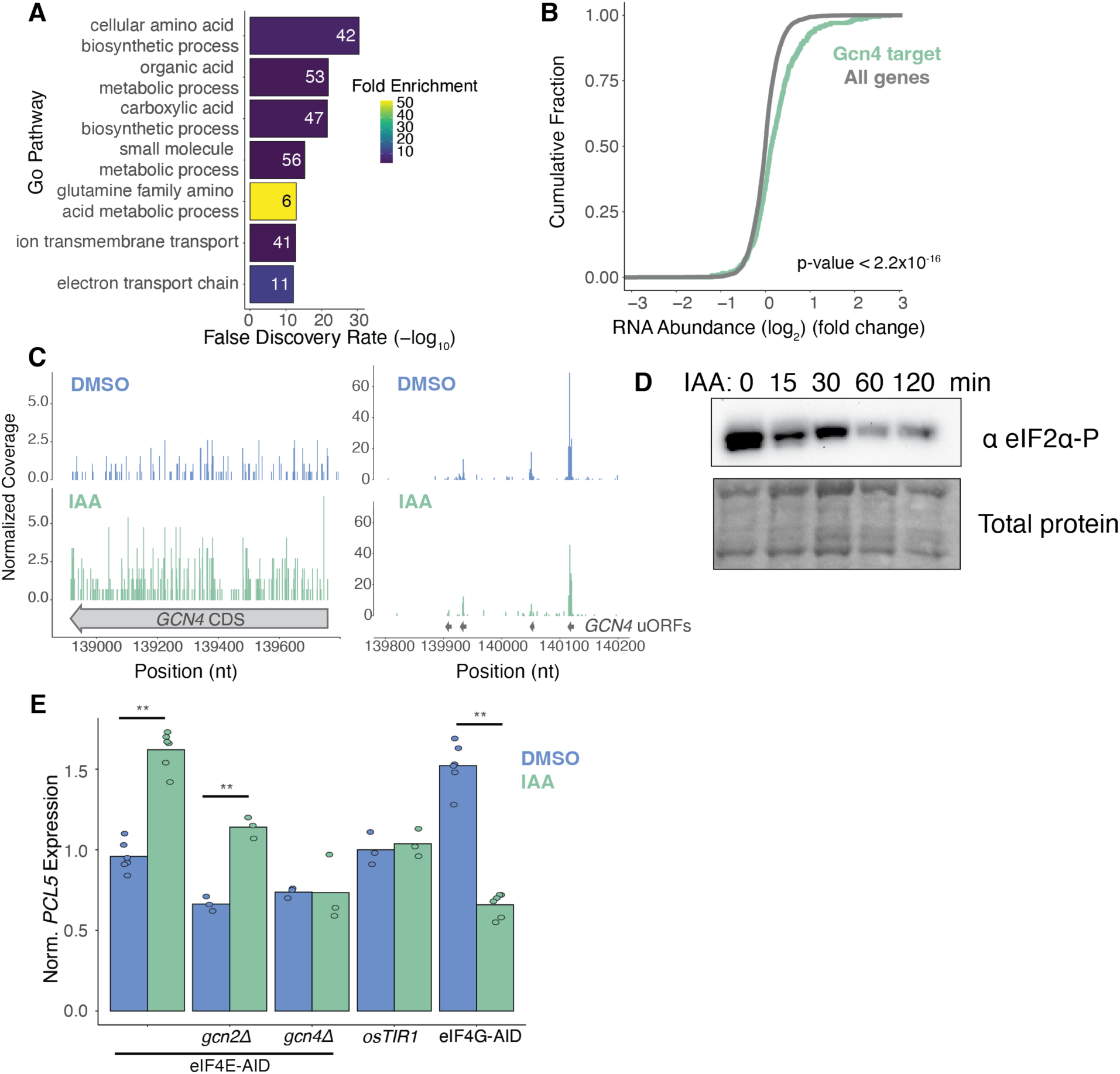
*GCN4* activation in response to eIF4E depletion. **(A)** GO analysis for genes which were significantly (adj. p-value < 0.05) up-regulated following eIF4E-depletion (8-hour treatment) in RNA-seq analysis. The most statistically significant entries were chosen and narrowed based on percentage of overlapping genes. **(B)** Empirical cumulative distribution function showing relationship between change in mRNA expression following eIF4E depletion for genes categorized as Gcn4 transcriptional targets. P-value was calculated using the Mann-Whitney U test. **(C)** Ribosome footprints (adjusted to A-site) over the GCN4 locus for 8-hour IAA and DMSO control cells. Read counts were scaled based on library size. **(D)** Western blot for eIF2α phosphorylation levels over the course of IAA depletion. **(E)** RT-qPCR of *PCL5* expression following 1-hour of IAA treatment. (**) represents p < 0.05 calculated by Student’s t test.

The translational induction of *GCN4* in response to amino acid starvation has been well characterized (Hinnebusch, 2005). During non-starvation conditions, the expression of *GCN4* is suppressed via repressive uORFs in the 5’UTR. This repression is relieved when the cell experiences amino acid starvation. When protein synthesis outstrips amino acid availability, uncharged tRNAs accumulate, causing the kinase Gcn2 to phosphorylate the translation initiation factor eIF2α. Phosphorylation of eIF2α reduces the availability of the active form of eIF2 bound to initiator tRNA, which allows reinitiating ribosomes in the *GCN4* 5’ UTR to bypass uORFs and reach the *GCN4* CDS, increasing Gcn4 protein levels (Dever et al., 1992). Surprisingly, although *GCN4* activation is canonically accompanied by an increase in eIF2a phosphorylation, we found that levels of eIF2α phosphorylation decreased over the course of eIF4E depletion (**Figure 3D**). Phosphorylation may decrease because reduced translation during eIF4E depletion reduced ribosomal collisions, another trigger for Gcn2 activation (Wu et al., 2020); in any case, the increased *GCN4* translation that we observed could not be explained by the Gcn2-eIF2α pathway.

To further dissect the genetic requirements for the novel mechanism of *GCN4* activation in eIF4E-depleted cells, we monitored the expression of the Gcn4 target gene Pho85 cyclin 5 (*PCL5)* in *gcn2Δ* and *gcn4Δ* cells (Shemer et al., 2002) (**Figure 3E**). We found that *PCL5* induction was Gcn4-dependent but Gcn2-independent, consistent with the lack of eIF2α phosphorylation (**Figure 3E**). To test whether translation deficiencies in general activated Gcn4 and thereby increased *PCL5* expression, we integrated AID tags at both paralogs of the translation initiation factor eIF4G, allowing us to conditionally deplete this essential translation factor. Depleting eIF4G significantly decreased *PCL5* expression, in contrast to the effects of eIF4E depletion (**Figure 3E**). We therefore concluded that the non-canonical Gcn4 activation following eIF4E depletion was not a result of general deficiencies in translation, but instead was a specific response to eIF4E-depletion.

### *PCL5* translation is regulated by uORFs and poly(A) tract in 5′ UTR

*PCL5* is a notable Gcn4 transcriptional target because it mediates a negative feedback circuit regulating Gcn4 activity (Shemer et al., 2002). The Pcl5 cyclin activates the cyclin-dependent kinase Pho85 to phosphorylate Gcn4, thereby marking it for degradation (Shemer et al., 2002). We reasoned that Gcn4 upregulation in eIF4E-depleted cells may reflect a breakdown in this negative-feedback loop. Curiously, when we evaluated the sequence of the 5′UTR of *PCL5*, we noted the presence of two potential uORFs with AUG start codons, followed by a startling stretch of 29 consecutive A bases— the longest such tract in the *S. cerevisiae* transcriptome (Vopálenský et al., 2019). Previous studies had also noted these potential AUG-start sites based on their conservation in yeast and the *PCL5* transcript structure, but they have heretofore been uncharacterized as regulators of *PCL5* translation (Cvijović et al., 2007; Zhang, 2005). The uORFs in the *PCL5* 5′UTR were reminiscent of the regulatory uORFs in the *GCN4* 5′UTR, which are responsible for its translational repression in un-starved conditions. The presence of *PCL5* uORFs could have important implications for Pcl5 protein synthesis during amino acid deprivation and therefore in the control of Gcn4 activity. By mapping ribosome occupancy across the 5′UTR of *PCL5*, we confirmed the translation of the two predicted AUG uORFs and, surprisingly, also observed translation of two uORFs beginning with non-cognate UUG codons (**Figure 4B**).

**Figure 4:**
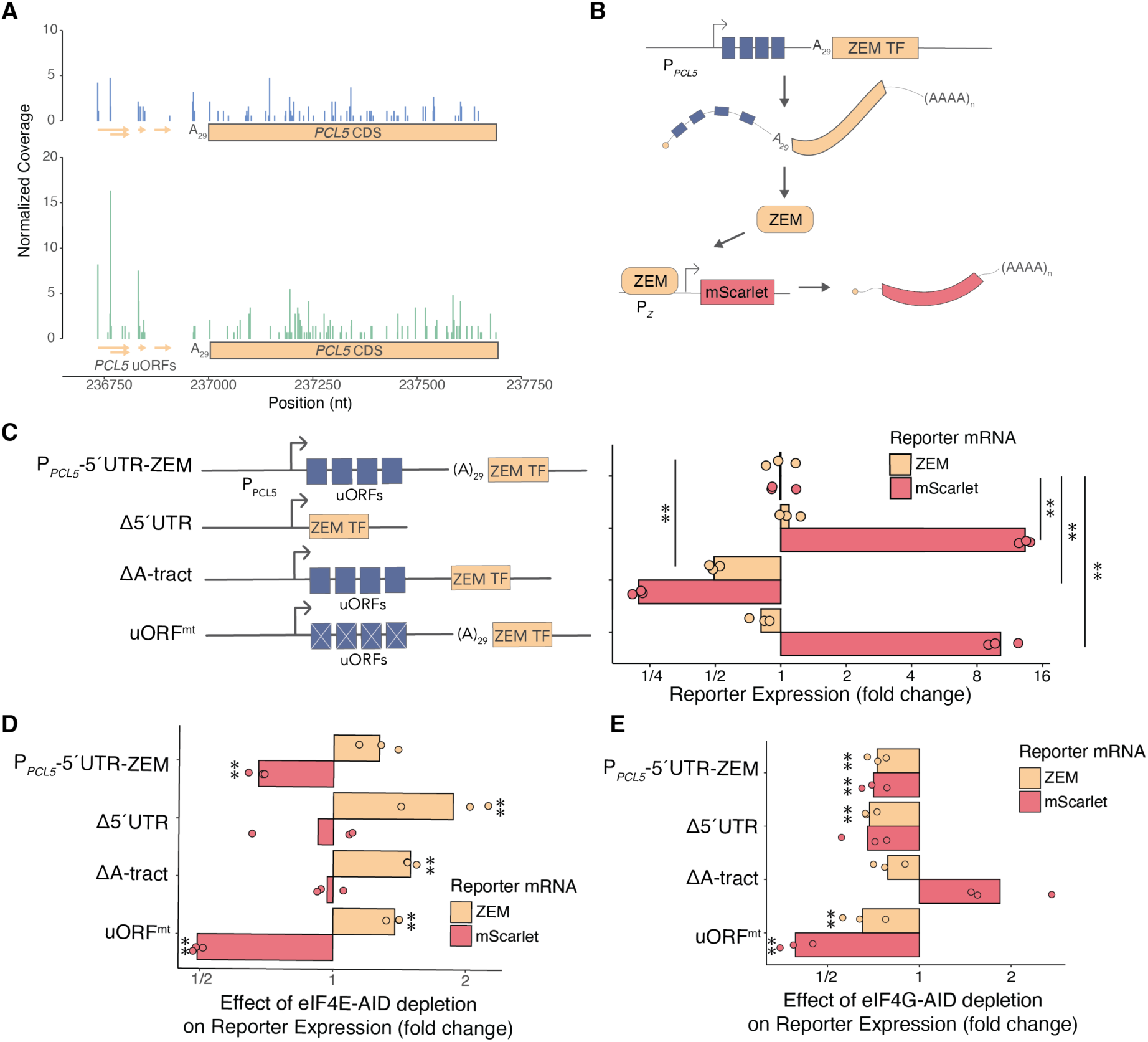
Translation regulation of *PCL5*. **(A)** Ribosome footprints (adjusted to A-site) over the *PCL5* locus for 8-hour IAA and DMSO control cells. Footprint counts were scaled based on library size and not adjusted for PCL5 abundance. **(B)** Schematic of *PCL5*-ZEM reporter assay. **(C)** RT-qPCR of ZEM transcript and mScarlet reporter expression of *PCL5* 5’UTR mutants normalized to 5’UTR^WT^ reporter. (**) represents p < 0.05 calculated by Student’s t test. **(D)** RT-qPCR of ZEM transcript and mScarlet reporter expression of *PCL5* 5’UTR mutants following eIF4E-AID depletion. Individual reporters were normalized to un-depleted control. (**) represents p < 0.05 calculated by Student’s t test. **(E)** Same as (D) except in eIF4G-AID depletion.

Interestingly, although *PCL5* was transcriptionally upregulated in response to eIF4E depletion (**Figure 3E**), we did not find a corresponding increase in ribosome occupancy over the *PCL5* ORF (**Figure 4A**). Instead, ribosome density in the *PCL5* 5’UTR was increased at all four uORFs in eIF4E-depleted cells, but most strikingly at the UUG uORF start codons (**Figure 4A**). In full, the translation efficiency of *PCL5* was reduced over 3-fold in response to eIF4E depletion (log_2_ fold-change = -1.77). We hypothesized that the distinctive regulatory features in the *PCL5* 5′UTR may have reduced its translation in eIF4E-depleted cells and sought to further characterize this regulation given the role of Pcl5 in repressing Gcn4 activity.

To further investigate how these unique *cis* features regulate *PCL5* translation, we designed fluorescent protein reporters fused to mutant versions of the *PCL5* 5′UTR, each testing the contribution of different 5′ UTR elements (**Figure S6A**). Unfortunately, most of reporters were weakly expressed and could not be distinguished from background fluorescence by flow cytometry (**Figure S6A**). We thus turned again to an indirect reporter system using the ZEM synthetic transcription factor, which amplifies signal from weakly expressed reporters (Aranda-Díaz et al., 2017) (**Figure 4B**). In this system, we measured changes to *PCL5-*reporter transcription and mRNA stability by tracking ZEM transcript levels (**Figure 4B**). Additionally, we measured *PCL5*-reporter translation indirectly, by measuring the abundance of a ZEM-driven transcriptional mScarlet reporter (**Figure 4B**). We generated four versions of this *PCL5-*ZEM reporter, each with the native *PCL5* promoter. We compared the full-length, wild type *PCL5* 5′UTR with three variants removing some or all of its distinctive *cis-*elements (**Figure 4C, S6B**). We then measured ZEM and mScarlet mRNA abundances to report on *PCL5* transcription and translation efficiencies, respectively. The *PCL5* uORFs suppressed translation of the ZEM transcription factor, consistent with the repressive effects typically seen from uORFs (**Figure 4C, S6B**). Furthermore, deletion of the poly-(A) tract reduced ZEM transcript levels by approximately 50%, and mScarlet transcript levels by 75%, relative to wild type (**Figure 4C, S6B**), suggesting that the long polyA tract in the *PCL5* 5′UTR enhanced its stability and translation.

Our previous findings indicated differential regulation of *PCL5* transcript levels in response to defects in translation initiation; Gcn4-dependent upregulation following eIF4E depletion and downregulation following eIF4G depletion (**Figure 3E**). To investigate whether *PCL5* translation was similarly affected by defects in translation initiation, we tracked changes to mRNA levels in these reporter constructs following depletion of eIF4E and eIF4G (**Figure 4D,E, S6C-F**). We observed an increase in ZEM-reporter transcript levels in all *PCL5-*ZEM reporters following eIF4E depletion, regardless of 5’UTR composition, likely mediated through the previously described *GCN4* response (**Figure 4D, S6C,D**). However, we found a decrease in mScarlet translation following eIF4E depletion in both the full-length 5’UTR reporter construct and the uORF mutation reporter.

In agreement with our measurements of endogenous *PCL5* transcript levels, eIF4G depletion consistently reduced ZEM-reporter transcript levels in all *PCL5-*ZEM reporters (**Figure 4E, S6E,F**). Strikingly, ZEM mRNA abundance was reduced even in the reporter that lacked the *PCL5* 5′UTR (**Figure 4E**, second line), suggesting that the regulation of *PCL5* in response to eIF4G depletion was transcriptional. Furthermore, we noted that expression of the reporter containing all four uORFs but lacking the polyA tract— the most repressive 5’UTR under wild-type conditions—was markedly less repressive following eIF4G-depletion. We therefore concluded that eIF4G may have played a role in suppressing translation in that context.

### Dissecting the genetic requirements for regulation of *PCL5* expression

Although the 5’UTR of *PCL5* contained sequence features that modulated its translation, we found that depleting eIF4E and eIF4G primarily affected its transcription rather than its translation. Thus, we investigated the potentially novel mechanisms regulating the translation of this factor by CiBER-seq profiling, using an indirect reporter of the same design as our *ARO10* reporter (**Figure 2**) (Muller et al., 2020). By fusing the ZEM transcription factor to the endogenous Pcl5 with a P2A self-cleaving peptide between the two proteins, we were able to capture regulators of *PCL5* transcription and translation using a transcriptional readout (**Figure 5A**). To parse the Gcn4-dependent transcriptional effects on *PCL5* from the translation regulation imparted through its 5’UTR, we performed parallel CiBER-seq analyses in a wild-type *GCN4* genetic background and one in which *GCN4* was deleted (*gcn4Δ)* (**Figure 5A**).

**Figure 5:**
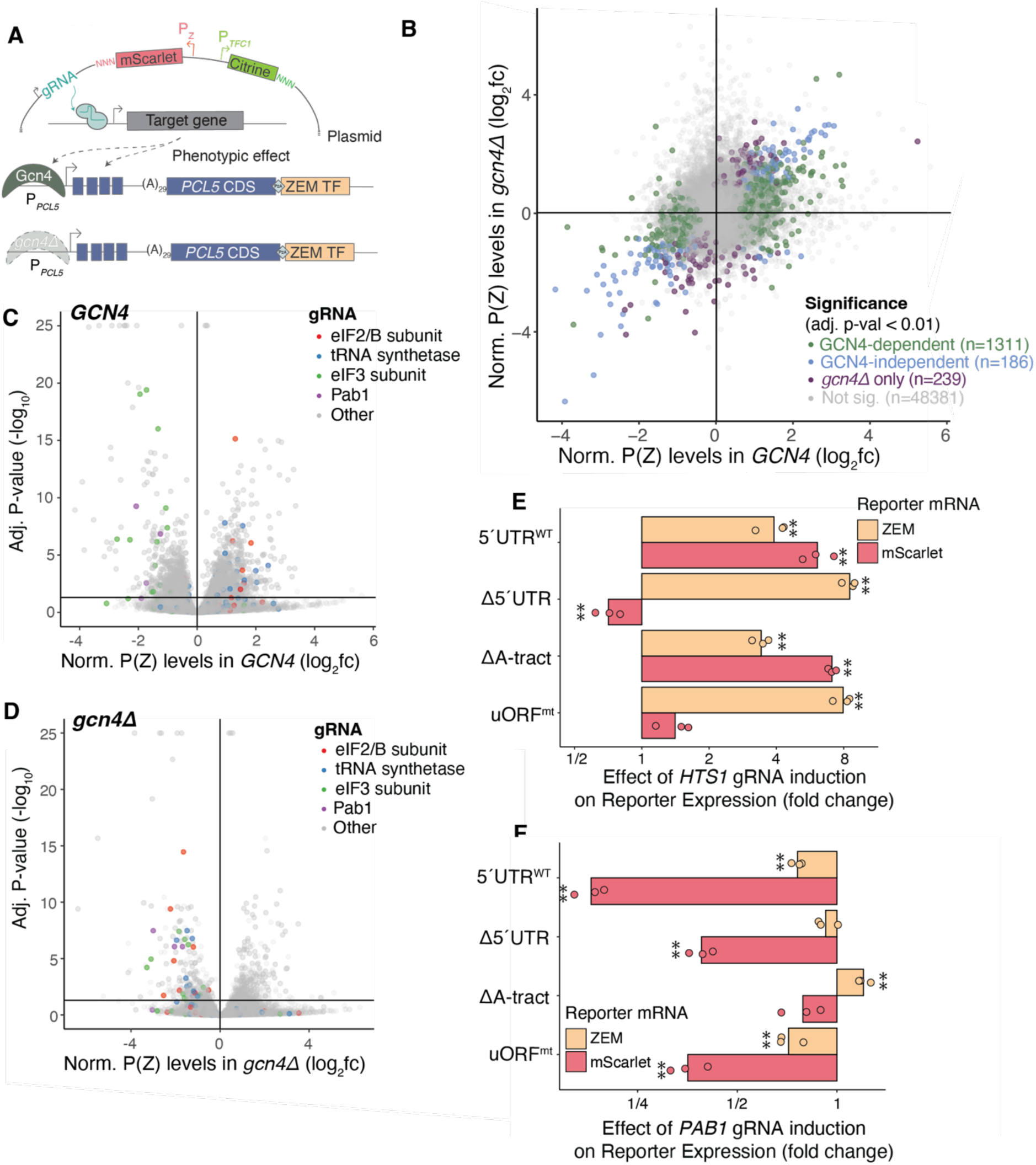
CiBER-seq genetic screen for regulators of *PCL5* expression. **(A)** Schematic of CiBER-seq screen design. **(B)** Genome-wide CiBER-seq screen results showing fold-change (log2) in P(Z) reporter abundance, relative to P(*TFC1*) reporter levels for each gRNA in *GCN4* and *gcn4Δ* backgrounds. Color indicates significance cutoff (adjusted p-value < 0.05). **(C)** Genome-wide CiBER-seq screen results showing fold-change (log2) in P(Z) reporter abundance, relative to P(*TFC1*) reporter levels for each gRNA in *GCN4* background. Line indicates significance cutoff (adjusted p-value < 0.05). Color represents gene category or identity. **(D)** Same as (C) in *gcn4Δ.* **(E)** RT-qPCR of ZEM transcript and mScarlet reporter expression of *PCL5* 5’UTR mutants following HTS1 gRNA induction. Individual reporters were normalized to uninduced control. (**) represents p < 0.05 calculated by Student’s t test. **(F)** Same as (E) except following *PAB1* gRNA induction.

Our screen identified thousands of guides that affected *PCL5* expression. Notably, the majority of these guides (1311/1780) had Gcn4-dependent effects (**Figure 5B**) and overlapped with previously published CiBER-seq analysis of *GCN4* itself (**Figure S7H,I**) (Muller et al., 2020). Many gRNAs known to activate the integrated stress response (ISR), such as those targeting tRNA synthetases and eIF2 or eIF2B subunits, induced *PCL5* expression in the *GCN4* wild type cells (**Figure 5C**). Interestingly, in *gcn4Δ* cells, these same gRNAs repressed *PCL5* expression (**Figure 5D**). This finding was surprising as we had anticipated similar regulation of *PCL5* and *GCN4* given the similarities between their 5’ UTRs. To further validate this finding, we tested a gRNA against the histidinyl-tRNA synthetase *HTS1*, whose knockdown is known to activate the ISR, in our *PCL5-ZEM* reporter system (**Figure 5E, S8B,E**). Consistent with results from our screen, knockdown of *HTS1* increased ZEM transcript levels and protein output (**Figure 5E, S8B,E**). Furthermore, we saw uORF-dependent regulation that maintained *PCL5* translation during ISR activation; reporters lacking the uORFs showed transcript-level increases that were not matched by increased protein output. (**Figure 5E, S8B,E**). This finding demonstrates for the first time that the translation of *PCL5* is regulated much like the translation of *GCN4* translation in response to eIF2a phosphorylation (Dever et al., 1992). These findings appeared to contradict our results from the *gcn4Δ* screen for regulators of *PCL5* translation (**Figure 5A,D**). To reconcile the two results, we hypothesize that there is a mechanism for suppressing *PCL5* expression when cells are unable to mount a sufficient Gcn4 response to ISR activation. Our screen recapitulated the *GCN4*-dependent induction of *PCL5* in response to loss of eIF4E and a contrasting, *GCN4*-independent reduction when eIF4G was depleted. We also observed system-dependent effects from knockdown of *PCL5* itself, the *GAL1*-derived reporter, and the *TFC1* normalizer (**Figure S7F,G**).

We were particularly interested in hits from our screen that regulated *PCL5* expression independently of Gcn4, as these may reveal mechanistic insight into translation control of *PCL5*. Knockdown of *PAB1*, which encodes the yeast poly-(A) binding protein, caused a *GCN4*-independent decrease in Pcl5 (**Figure 5D**)—a striking observation in light of the 29 base poly-(A) tract in the *PCL5* 5’ UTR, which is long enough to bind Pab1 (Sachs et al., 1987). Indeed, we tested the effects of *PAB1* knockdown on mutant *PCL5* reporters and found that removal of the poly-(A) tract greatly attenuated the strong translational repression seen in the wild-type version (**Figure 5F, S8D,G**). We likewise observed that guides targeting subunits of the translation initiation factor eIF3 reduced *PCL5* reporter translation even in the absense of *GCN4*. As eIF3 has known roles in ribosome recycling and re-initiation, particularly in the context of the uORFs in the *GCN4* 5’UTR (Sonenberg and Hinnebusch, 2009), we confirmed that knockdown of eIF3 subunit gene *NIP1* reduced reporter mRNA and translation levels in a uORF-dependent manner (**Figure S8A,C,F**). These results, which matched our CiBER-seq data, suggested that eIF3 supports translation reinitiation downstream of *PCL5* uORFs as well. Together, our findings highlighted the complexity of mechanisms governing *PCL5* expression and suggested that regulation of *PCL5* serves as an integration point of different signaling pathways to sculpt the Gcn4 response.

## Discussion

We showed that acute depletion of the essential cap-binding protein eIF4E had surprisingly modest effects on cell growth and protein synthesis. Although eIF4E levels were quickly reduced below 10% of starting abundance, we observed a relatively uniform 50% reduction in overall translation. We did note more nuanced changes to mRNA abundance, including a strong reduction in the expression of long-lived transcripts, likely reflecting more rapid turnover (Chan et al., 2018). There are conflicting data on whether the eIF4E-mRNA interaction is rate-limiting for cap-dependent translation initiation (Firczuk et al., 2013; von der Haar et al., 2004). Furthermore, eIF4E concentration exceeds that of mRNA several-fold, while eIF4G has roughly matched stoichiometry with mRNA. Notably, this implies the majority of eIF4E is not in complex with eIF4G and may be stabilizing or promoting translation of specific transcripts (Firczuk et al., 2013; Krause et al., 2022). In contrast to our results, prior work reported that transcriptional titration of eIF4E led to proportional decreases in protein synthesis and growth rates (Firczuk et al., 2013; von der Haar et al., 2004). In mice, however, eIF4E haploinsufficiency does not cause bulk translation defects (Truitt et al., 2015), suggesting that the relative insensitivity to loss of cap-binding protein might be a conserved aspect of translation biology (O’Leary et al., 2013; von der Haar and McCarthy, 2002).

We further showed that protein synthesis recovered after only four hours, although eIF4E levels remained extremely low, suggesting that cells robustly adapted to this condition. Following prolonged eIF4E depletion, cells exhibited more substantial changes to translation than seen at earlier timepoints. These long-term translation changes weakened the bias towards more efficient translation of short transcripts. This trend is intriguing given that efficient translation of short transcripts is attributed to the formation of a closed-loop structure, wherein eIF4G bridges between eIF4E at the 5′ end of an mRNA and the poly(A)-binding protein, Pab1, at its 3′ end (Amrani et al., 2008; Çetin and O’Leary, 2022; Costello et al., 2015; Thompson and Gilbert, 2017). Depletion of eIF4E would abolish closed-loop formation, explaining the reduced the translation efficiency of short mRNAs and depressed the length bias of translation efficiency.

The strongest gene-specific effects of eIF4E depletion arose as secondary effects of reduced protein biosynthesis on amino acid pools. The transcription factor Aro80 responded to changes in translational status by activating a strong catabolic response through Ehrlich pathway enzymes Aro9 and Aro10, a gene expression program observed under different starvation stress conditions (Staschke et al., 2010; Xia et al., 2022; Zou et al., 2020). In the context of eIF4E depletion, this catabolic response appears maladaptive as cells require ongoing aromatic amino acid biosynthesis or supplementation to grow. It may play other roles, however—aromatic alcohols produced by this pathway serve as quorum sensing molecule in *S. cerevisiae* (Chen and Fink, 2006; Zhang et al., 2021).

Paradoxically, reductions in eIF4E activity also caused translational activation of *GCN4* and thereby induced a transcriptional response that favored amino acid anabolic processes. Surprisingly, this activation of *GCN4* occurred independently of Gcn2 and without eIF2α phosphorylation, indicating the involvement of a non-canonical mechanism. Fusel alcohols produced by amino acid catabolism can regulate *GCN4* translation through the inhibition of eIF2B, acting downstream of eIF2α phosphorylation, offering one possible explanation (Ashe, 2001; Hazelwood et al., 2008). Beyond this possibility, we have recently described a range of genetic perturbations to translation that lead to Gcn2-independent *GCN4* induction (Muller et al., 2020). In the case of eIF4E depletion, reduced recruitment of ribosomal 40S subunits for translation could increase the availability of free 40S subunits and thereby depress the ratio of eIF2/40S, similar to the *GCN4* activation caused by large subunit biogenesis (Steffen et al., 2008).

The translational repression of *PCL5*, a negative regulator of Gcn4, may have further imbalanced amino acid pools following eIF4E depletion. Pcl5 promotes Gcn4 degradation. Because *PCL5* expression is induced by Gcn4, this regulated degradation serves as a feedback mechanism that controls Gcn4 activity (Shemer et al., 2002). Feedback control requires that Pcl5 protein synthesis is proportional with its Gcn4-driven transcription, but maintaining the correspondence between mRNA levels and protein production poses a challenge when translation is impaired due to intrinsic limitations, inhibition by eIF2α phosphorylation, or other factors—exactly the conditions when Gcn4 regulation is needed. The requirement for uniform translation under varying conditions may explain why *PCL5* contains uORFs similar to those that control *GCN4* translation. Our findings highlight the significance of these characteristics as a point of regulation for *PCL5* that shapes the integrated stress response in yeast.

The post-transcriptional control of *PCL5* has distinctive characteristics, however, which may be mediated through the poly(A) tract in its 5′ UTR. Our finding that the poly(A) tract and Pab1 stabilize the *PCL5* transcript is reminiscent of previous work that found internal Pab1-tethering recruited the termination factor eRF3 and stabilized nonsense-containing mRNAs (Amrani et al., 2004). Additionally, *PCL5* translation regulation may integrate additional signals about cellular environment, such as heat shock, a condition in which Pab1 has been shown to phase separate (Riback et al., 2017). While poly(A) tracts have previously been linked to cap-independent translation of internal ribosome entry sites in yeast (Gilbert et al., 2007), translation of *PCL5* appears to be cap-dependent; it is repressed by uORFs, which affect cap-dependent scanning, and it is reduced upon depletion of cap-binding eIF4E (**Figure 4**).

Amino acid homeostasis is crucial for cell growth and survival. The tight regulation of amino acid pools involves a complex interplay between amino acid uptake, biosynthesis, incorporation into protein, and catabolism. Hundreds of genes are regulated response to changes in amino acid availability. Our study has identified the cap-binding protein eIF4E as a central coordinator of both biosynthetic and catabolic gene expression **(Figure 6**). Depletion of eIF4E led to upregulation of both opposing metabolic processes, resulting in an excess of aromatic amino acids and a discordant upregulation of amino acid biosynthesis genes, potentially indicative of gratuitous or futile amino acid metabolism. Intriguingly, mammalian cells couple amino acid sensing with protein synthesis through cap-centric regulation as well, albeit through a distinct mechanism. Indeed, dysregulation of eIF4E and metabolism are hallmarks of cancer, emphasizing the importance of understanding the underlying mechanisms to develop effective therapeutic strategies.

**Figure 6:**
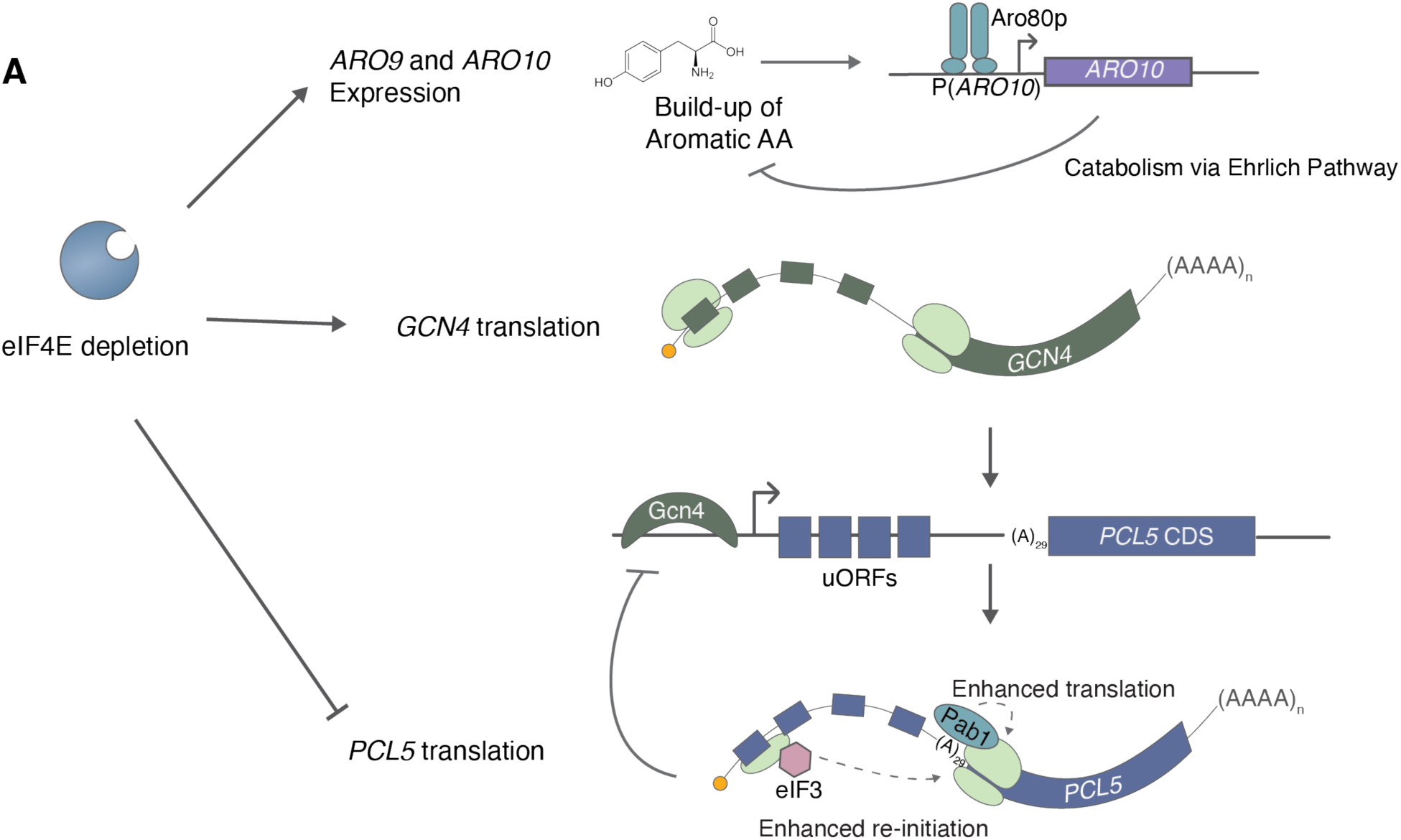
Model of the role of eIF4E in maintenance of amino acid homeostasis. **(A)** Model emphasizing the effects of eIF4E depletion on dysregulation of amino acid metabolism gene expression and mechanisms of *PCL5* translational regulation.

## Materials and Methods

### Yeast strains

Strains used in this study are listed in Table S1. All strains were derived from the S. cerevisiae BY4742 using standard genetic techniques and CRISPR-Cas9 technology (Lee et al., 2015). C-terminal mAID-3xFlag tags (cdc33-mAID-3xFlag, tif4631-mAID-3xFlag) were generated by amplifying the SG_linker-mAID-3xFlag sequence from pNTI433 with primers that included 40 bp of sequence identity with either side of the insertion, then integrated using CRISPR-Cas9 technology as described by (Brothers and Rine, 2019). C-terminal mAID-3xV5 tag (tif4632-mAID-3xV5), and P2A-ZEM tags (aro10-P2A-ZEM and pcl5-P2A-ZEM) were constructed in the same manner by amplifying from synthetic DNA gene block (Integrated DNA Technologies) PD552 and pNTI730, respectively. Gene deletions were generated using one-step replacement with marker cassettes, more specifically by PCR amplification of the hygromycin resistance cassette from pNTI730 with flanking sequence homology to the 5’ and 3’ ends of target gene coding sequence (Goldstein and McCusker, 1999; Hentges et al., 2005). *PCL5*-ZEM reporters and dCas9-TetR were integrated using NotI-linearized vectors from the EasyClone 2.0 toolkit for yeast genomic integration (Stovicek et al., 2015). Plasmids and primers used in strain construction are listed in Table S2 and S3, respectively.

### Plasmids

Plasmids used in this study are listed in Table S2. For CRISPR-Cas9 editing, guide RNA target and nontarget strands were integrated into single guide RNA dropout-Cas9 expression plasmid (pJR3429, a gift from the Rine lab) as described in (Brothers and Rine, 2019). Annealing oligos for gRNA insertion are listed in Table S3. The PCL5-ZEM reporter plamids (PDp86-89) were made using standard Gibson cloning into pCfB2337 (Gibson et al., 2009; Stovicek et al., 2015).The PCL5-mCherry reporter plasmids (PDp74-78) were made using standard Gibson cloning into pRS315 LEU2 CEN/ARS (a gift from the Rine lab). P(*Z*)-mScarlet and P(*TFC1*)-citrine reporter plasmids with individual guide RNAs were made through standard Gibson cloning of annealed oligos into a single isolate of the plasmid library.

### HPG assay

Yeast cells were inoculated in a custom turbidostat (McGeachy et al., 2019) and maintained in SCD -Met media at a density corresponding to OD_600_ of 0.6. After growth rate reached a steady state, indole-3-acetic acid (IAA) (Sigma-Aldrich #12886) was injected into the growth chamber and media reservoir to achieve a final concentration of 500 µM. After indicated duration of eIF4E depletion, 5 mL of cells were collected for western blot analysis and HPG nascent peptide labeling.

For HPG nascent peptide labeling assay, cells were back-diluted to OD_600_ of 0.3 in SCD -Met with 500 µM IAA and 50 µM L-Homopropargylglycine (HPG) (Thermo Fisher Scientific #C10186). Nascent peptide labeling was conducted for 2 hours at 30°C. For cycloheximide control samples, 50 µg/mL cycloheximide (Sigma-Aldrich #C4859) was added concurrently with HPG. Cells were harvested by centrifugation and fixed at room temperature while nutating in 4% paraformaldehyde (Electron Microscopy Sciences #15710). Cells were washed with PBS with 2mM EDTA and permeabilized at 45°C with 2% w/v sodium lauroyl sarcosine (Bioworld #41930024-3) in PBS for 15 minutes (Abraham and Bhat, 2008). Click-iT^TM^ chemistry was performed per manufacturer’s recommendations (Thermo Fisher Scientific #C10428) to label HPG with Alexa Fluor™ 488 azide. Fluorescence was measured by flow cytometry analysis on a BD LSR Fortessa X20 with excitation by the 488 nm blue laser, captured on the FITC channel.

### Polysome profiling

Cells were grown to mid-log phase (typically 100 mL cultures at OD_600_ 0.6 – 1.0) in YPD, harvested by filtration with 0.45 µM Whatman cellulose nitrate membrane filters (GE Life Sciences #7184-004). 100 µg/mL cycloheximide was added to the culture immediately before filtration. Samples were lysed by cryogrinding with the MM400 Mixer mill (Retsch #20.745.0001) in polysome buffer (20 mM Tris-HCl pH 7.4, 150 mM NaCl, 5 mM MgCl_2_, 1 mM DTT, 100 µg/mL cycloheximide), followed by centrifugation at 10,000 g for 10 min at 4°C. The supernatant was collected and stored at -80°C.

Sucrose gradients were prepared using a Gradient Master (BioComp Instruments, Fredericton, Canada). Normalized lysates (200 µL) were loaded onto a 10-50% sucrose gradient prepared in polysome buffer. Gradients were centrifuged at 36,000 rpm for 4 hours at 4°C in a Beckman SW41Ti rotor. The gradient run through the Gradient Master, and the absorbance at 254 nm was monitored continuously using a BioRad EM-1 Econo UV Monitor.

### Western blotting

Cells were grown to mid-log phase (typically 5 mL cultures at OD_600_ 0.6 – 1.0) in YPD, harvested by centrifugation, and washed with 20% TCA (Cox et al., 1997). The pellet was flash frozen in liquid nitrogen and stored at -80°C until further use. For lysis and extraction, the pellet was thawed on ice and resuspended in 20% TCA (Sigma-Aldrich #T6399) and glass beads (Sigma-Aldrich #Z250465) The sample was then lysed by vortexing and centrifuged at 4°C, 20,000xg for 10 minutes. The pellet was washed with ice-cold 100% ethanol and centrifuged again. The final pellet was resuspended in Tris pH 8 buffer.

Equal amounts of lysate were denatured for 10 minutes at 80°C and loaded on a 4 to 12% polyacrylamide Bis-Tris gel (Thermo Scientific #NW04120BOX). The gel was run at 120V for approximately 80 minutes in MOPS buffer. Protein was then transferred to a nitrocellulose membrane (Thermo Scientific #88018) according to manufacturer’s guidelines. Membranes were blocked for 1 hour in TBST with 0.5% milk. Primary antibodies were incubated with the membrane overnight at 4°C while shaking and secondary antibodies were incubated for 1 hour at room temperature. Membranes were washed three times by shaking for 5 minutes in TBST following primary and secondary incubations. All blots were developed with Pierce ECL Western Blotting Substrate (Thermo Scientific #32209) and imaged by a FluorChem R imaging system (ProteinSimple). The antibodies and corresponding dilution factors used in this study include rabbit anti-Flag (CST #2368S) (1:1000), rabbit anti-V5 (CST #13202) (1:1000), rabbit anti-HA (CST #3724) (1:1000), mouse anti-GAPDH (Proteintech #60004-1-IG) (1:5000), rabbit anti-Phospho-eIF2α(Ser51) (CST #3398S) (1:1000), HRP-conjugated anti-rabbit IgG (CST #7074) (1:5000).

### Ribosome profiling and RNA-sequencing

Cells were treated with 500 µM indole-3-acetic acid (IAA) (Sigma-Aldrich #12886) or DMSO for either 1 or 8 hours and grown to mid-log phase (typically 150 mL cultures at OD_600_ 0.6 – 1.0) in YPD. Yeast cells were harvested by filtration with 0.45 µM Whatman cellulose nitrate membrane filters (GE Life Sciences #7184-004). Ribosome profiling was conducted as detailed in (McGlincy and Ingolia, 2017) for 1-hour eIF4E depletion samples and corresponding DMSO controls. Adjustments were made to (McGlincy and Ingolia, 2017) for the 8-hour eIF4E depletion samples and their corresponding DMSO control samples as follows. Reverse transcription was performed with primer PD1131. rRNA depletion was achieved via subtractive hybridization was performed using complementary oligos PD1043-1046 as described in (Ingolia et al., 2012). Circularization was performed with CircLigase I (Epicentre #CL4111K) instead of CircLigase II, following the manufacturer’s protocol recommendations.

Total RNA for RNA-sequencing was isolated from cell lysates using acid phenol extraction (Ares, 2012). For 1-hour eIF4E depletion samples and corresponding DMSO controls, poly(A) enrichment was performed using Dynabeads^TM^ oligo(dT)_25_ (Thermo Scientific #61002) according to the manufacturer instructions. For 8-hour eIF4E depletion samples and corresponding DMSO controls, rRNA depletion was performed using QIAseq FastSelect -rRNA yeast kit (Qiagen #334215) according to the manufacturer instructions. All libraries were generated with NEBNext Ultra II Directional RNA library prep kit (NEB #E7760).

Reads from ribosome profiling were processed as detailed in (McGlincy and Ingolia, 2017). Ribosome profiling and RNA-seq reads were aligned using STAR (Dobin et al., 2013) to the S288C reference genome R64.1. Differential expression analysis was performed with DESeq2 (Love et al., 2014).

### Plasmid library generation and transformation

The divergent *P(Z)-*mScarlet and *P(TFC1)-*citrine plasmid libraries were generated as detailed in (Muller et al., 2022, 2020) with the following modifications. The divergent promoter inserts for Gibson assembly (NEB #E2621L) were generated by digesting PDp82 with SalI and AvrII. These libraries were transformed into ElectroMax DH10B cells (Thermo Scientific, #18290015) by electroporation according to manufacturer’s protocol. Serial dilutions of transformations were plated to ensure sufficient library diversity (>50x coverage of 240,000 unique barcodes).

Plasmid libraries were transformed into yeast using standard lithium acetate transformation (Gietz and Schiestl, 2007). Serial dilutions of transformations were plated to ensure sufficient library diversity (>5x coverage of 240,000 unique barcodes).

### CiBER-seq

Yeast populations transformed with plasmid libraries were inoculated in a custom turbidostat (McGeachy et al., 2019) and maintained in SCD -URA media with 10 nM beta-estradiol at a density corresponding to OD_600_ of 0.6. After a period of six doublings (9 hours), a sample of 50 mL was taken prior to the induction of gRNAs. The induction was carried out by adding anyhydrotetracycline (Abcam Biochemicals #ab145350) to both the growth chamber and the turbidostat media reservoir, in order to achieve a final concentration of 250 ng/ml. Following another six doublings, which took another 9 hours, a post-induction sample of 50 mL was collected.

RNA extractions and barcode library preparations were performed as previously described (Muller et al., 2022, 2020). The cDNA products were amplified with PD780 and RM810 for P(*Z*)-mScarlet libraries, and RM810 and RM511 for P(*TFC1*)-citrine libraries. Sequencing data was first subjected to processing using Cutadapt (Martin, 2011) to trim adapter sequences and deconvolve multiplexed libraries based on embedded nucleotide indices. Trimmed barcodes were then counted as described in (Muller et al., 2020). Barcodes with fewer than 32 counts in at least one of the replicates of the pre-induction samples were excluded from the analysis. The remaining barcodes were evaluated by differential activity analysis using mpralm (Ashuach et al., 2019).

### RT-qPCR

For ZEM RT-qPCR assays, cells were back-diluted from saturation and grown for 9 hours in SCD -URA with 10 nM beta-estradiol and 250 ng/ml anyhydrotetracycline (Abcam Biochemicals #ab145350) when applicable. For eIF4E depletion RT-qPCR assays, cells were back-diluted from saturation and grown for 4 hours in YPD with 500 µM IAA (Sigma-Aldrich #12886) or DMSO. All cells were harvested by centrifugation from mid-log phase. RNA was extract using either acid phenol extraction (Ares, 2012) or Direct-zol RNA miniprep kit (Zymo Research #R2053). Samples extracted with Direct-zol RNA miniprep kit were disrupted by vortexing with glass beads (Sigma-Aldrich #Z250465) for 5 minutes in TRI reagent. RNA samples were treated with DNase I (Thermo Scientific #EN0521) for 1 hour at 37°C, followed by RNA clean up with RNA Clean & Concentrator kit (Zymo Research R1015).

Complementary DNA was synthesized using the SuperScript II reverse transcriptase (Invitrogen #18064014) and random primers according to the manufacturer’s protocol. Quantitative PCR was conducted using the DyNAmo HS SYBR Green kit (Thermo Scientific #F410L) and the primers specified in Table S1. The procedure was carried out on an Mx3000P machine (Stratagene, La Jolla, CA), and standard curves were created using a tenfold dilution sequence of one of the samples that had been prepared.

### β-estradiol titrations

Cells were back-diluted from saturation and grown in SCD -URA with various β-estradiol (Sigma-Aldrich #E8875) concentrations for 8 hours to mid-log phase. Cells were fixed in 4% PFA for 15 minutes, washed and resuspended in PBS + 2 mM EDTA. Fluorescence was measured a BD LSR Fortessa X20 with excitation by the 561 yellow-green laser and captured on the PE-TexasRed channel.

### Growth assays

Cells were grown to mid-log phase (typically 5 mL cultures at OD_600_ 0.6 – 1.0 in YPD unless otherwise indicated), and back-diluted to OD_600_ 0.025 in triplicate. Absorbance measurements at 595nm were measured every 15 min using a 96-well plate reader (Tecan SPARK Multimode Microplate Reader). Growth rates were fit using the R package ‘growthcurver’ (Sprouffske and Wagner, 2016).

## Acknowledgments

We are grateful to the members of the Ingolia laboratory for helpful discussions in the planning of this work. We give special thanks to Eliana Bondra and Joe Lobel for their keen editing. This work relied on the Vincent J Coates Genomics Sequencing Laboratory at UC Berkeley. This work was supported by the National Institutes of Health (www.nih.gov), grants DP2 CA195768 and R01 GM130996 (N.T.I.) and shared instrumentation grant S10 OD018174. The funders had no role in study design, data collection and analysis, decision to publish, or preparation of the manuscript.

## Data Availability

All sequencing files are available on NCBI GEO database (accession number GSE231759).

---

**Figure S1:**
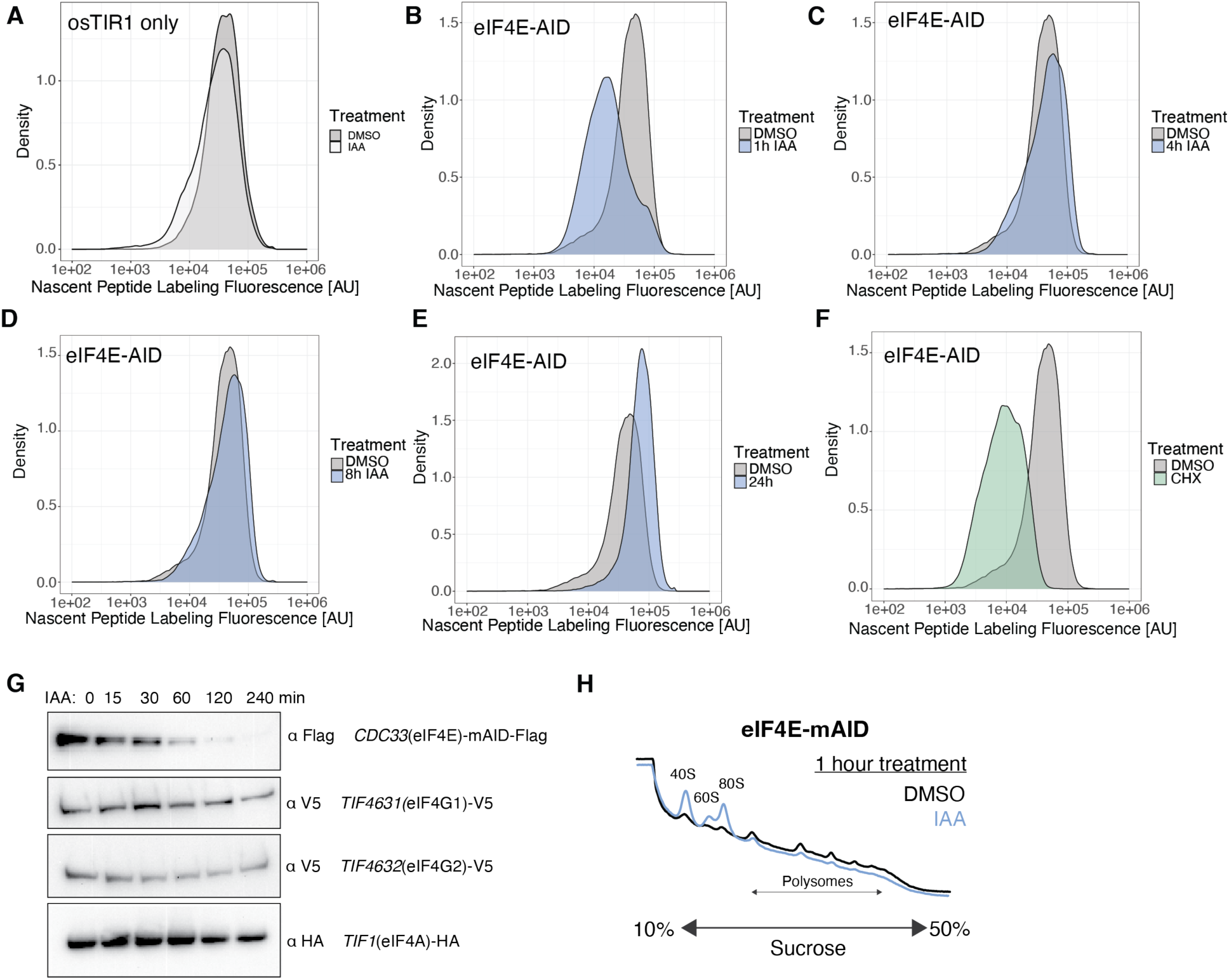
Impact of eIF4E depletion on bulk protein synthesis. **(A)** Bulk translation measured by nascent peptide metabolic labeling in *osTIR1* only expressing cells. Indole-3-acetic acid (IAA) or DMSO was added for 1 hour and maintained during a 2-hour labeling period with L-Homopropargylglycine (HPG). Fluorescence intensity of Alexa Fluor^TM^ 488 (HPG) signal was measured with flow cytometry. **(B)** Same as in (A) for eIF4E-AID cells. **(C)** Same as in (B) except IAA was added for 4 hours before labeling period. **(D)** Same as in (B) except IAA was added for 8 hours before labeling period. **(E)** Same as in (B) except IAA was added for 24 hours before labeling period. **(F)** Same as in (A) except cycloheximide (CHX) was added concurrently with HPG. **(G)** Western blot analysis for translation initiation expression levels over the course of eIF4E-AID-flag depletion. **(H)** Polysome profiles eIF4E-AID cells treated with IAA or DMSO for 1 hour.

**Figure S2:**
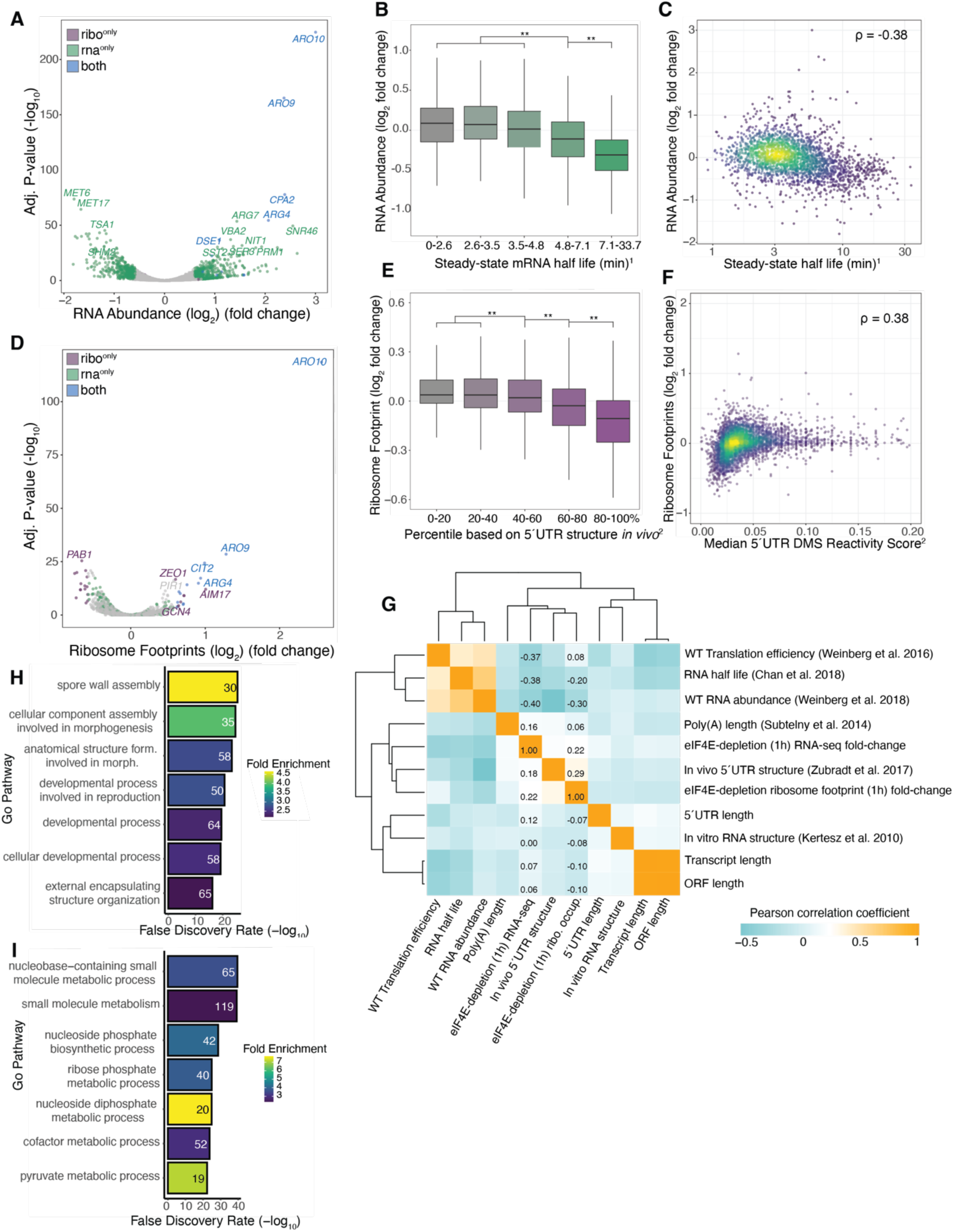
Transcript-level sensitivities after 1 hour of eIF4E depletion. **(A)** Differential expression after 1 hour of eIF4E depletion measured by RNA-seq. IAA-treated cells are compared with DMSO-treated controls. Color represents significant (adjusted p-value < 0.05) and substantial (absolute fold-change (log2) > 0.58) changes for RNA-seq and ribosome profiling measurements. **(B)** Box plots of RNA abundance fold change (log2) after 1 hour of eIF4E depletion for transcripts grouped based on steady-state mRNA half-life (Chan et al., 2018). (**) indicates p < 0.05; one-way ANOVA test followed by Tukey’s HSD test. **(C)** Scatterplot of steady-state mRNA half life (Chan et al., 2018) and fold change (log2) after 1 hour of eIF4E depletion measured by RNA-seq. Color represents point density. Correlation coefficient (Spearman’s) calculated between log2 fold changes of RNA abundance and half-life (min). **(D)** Same as (A) except changes in ribosome occupancy measured by ribosome profiling. **(E)** Box plots of ribosome occupancy fold change (log2) after 1 hour of eIF4E depletion for transcripts grouped based on *in vivo* 5’UTR structure (median DMS reactivity) from DMS-MaPseq data (Zubradt et al., 2017). (**) indicates p < 0.05; one-way ANOVA test followed by Tukey’s HSD test. **(F)** Scatterplot of DMS reactivity scores over the 5’UTR of transcripts based on DMS-MaPseq data (Zubradt et al., 2017) and fold change (log2) after 1 hour of eIF4E depletion measured by ribosome profiling. Color represents point density. Correlation coefficient (Spearman’s) calculated between log2 fold changes of RPFs and DMS reactivity score. **(G)** Spearman’s correlation coefficients of pairwise comparisons across mRNA parameters including translation efficiency(Weinberg et al., 2016), stability (Chan et al., 2018), expression, poly(A) length (Subtelny et al., 2014), 5’UTR structure (Kertesz et al., 2010; Zubradt et al., 2017), and transcript length. Datasets were hierarchically clustered based on Euclidian distances. Color represents magnitude of correlation coefficient. **(H)** GO analysis for genes which were significantly (adj. p-value < 0.05) up-regulated following eIF4E-depletion (1-hour treatment) in RNA-seq analysis. The most statistically significant entries were chosen and narrowed based on percentage of overlapping genes. **(I)** Same as in (H) for down-regulated genes.

**Figure S3:**
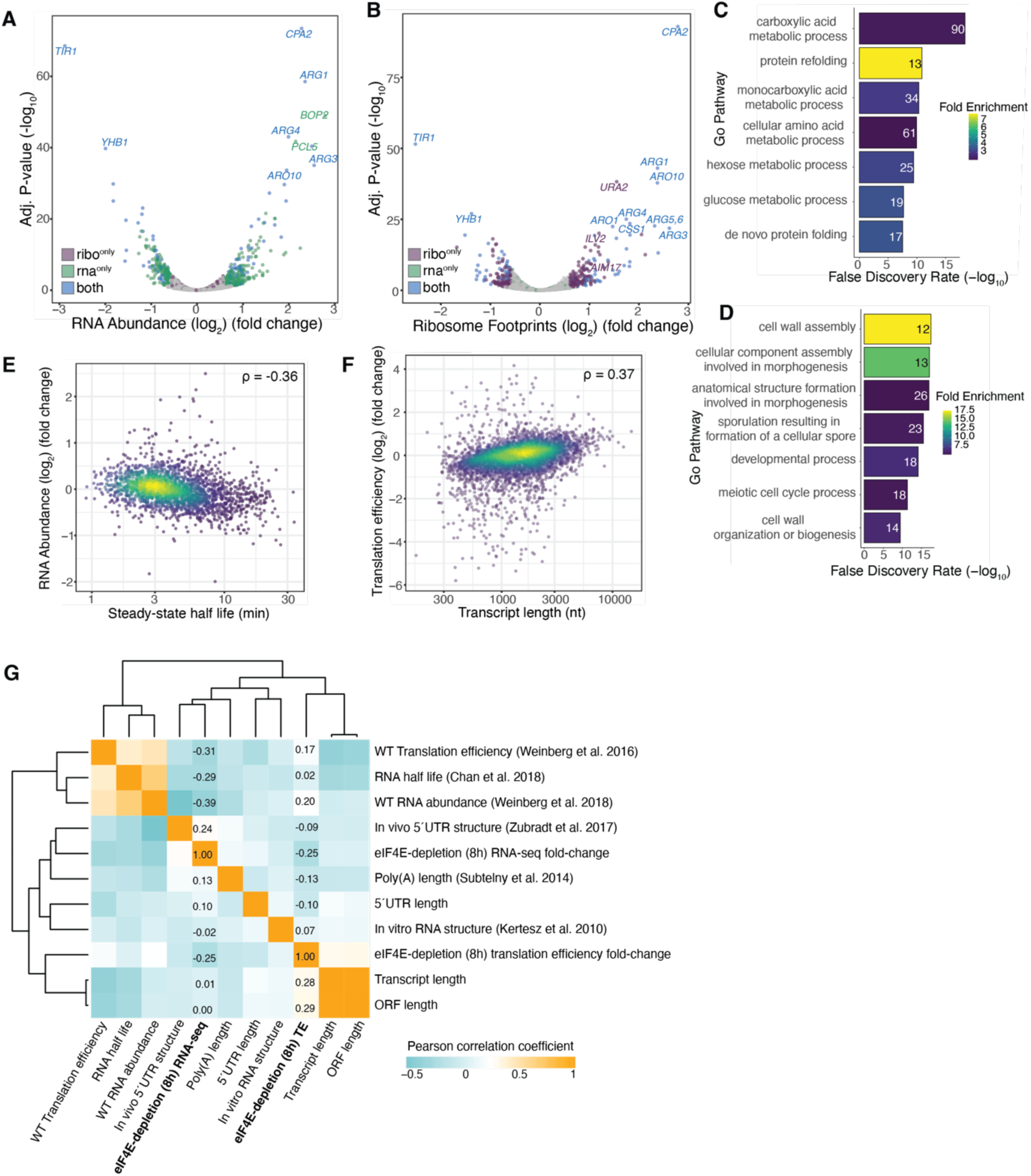
Transcript-level sensitivities after 8 hour of eIF4E depletion. **(A)** Differential expression after 8 hour of eIF4E depletion measured by RNA-seq. IAA-treated cells are compared with DMSO-treated controls. Color represents significant (adjusted p-value < 0.05) and substantial (absolute fold-change (log2) > 0.58) changes for RNA-seq and ribosome profiling measurements**. (B)** Same as (A) except changes in ribosome occupancy measured by ribosome profiling. **(C)** GO analysis for genes which were significantly (adj. p-value < 0.05) up-regulated translation efficiency following eIF4E-depletion (1-hour treatment). The most statistically significant entries were chosen and narrowed based on percentage of overlapping genes. **(D)** Same as in (C) for genes downregulated in translation efficiency. **(E)** Scatterplot of steady-state mRNA half-life (Chan et al., 2018) and fold change (log2) after 8 hour of eIF4E depletion measured by RNA-seq. Color represents point density. Correlation coefficient (Spearman’s) calculated between log2 fold changes of RNA abundance and half-life (min). **(F)** Scatterplot of transcript length and fold change (log2) in translation efficiency (TE) after 8 hours of eIF4E depletion. Color represents point density. Correlation coefficient (Spearman’s) calculated between log2 fold changes of TE and transcript length. **(G)** Spearman’s correlation coefficients of pairwise comparisons across mRNA parameters including translation efficiency (Weinberg et al., 2016), stability (Chan et al., 2018), expression, poly(A) length (Subtelny et al., 2014), 5’UTR structure (Kertesz et al., 2010; Zubradt et al., 2017), and transcript length. Datasets were hierarchically clustered based on Euclidian distances. Color represents magnitude of correlation coefficient.

**Figure S4:**
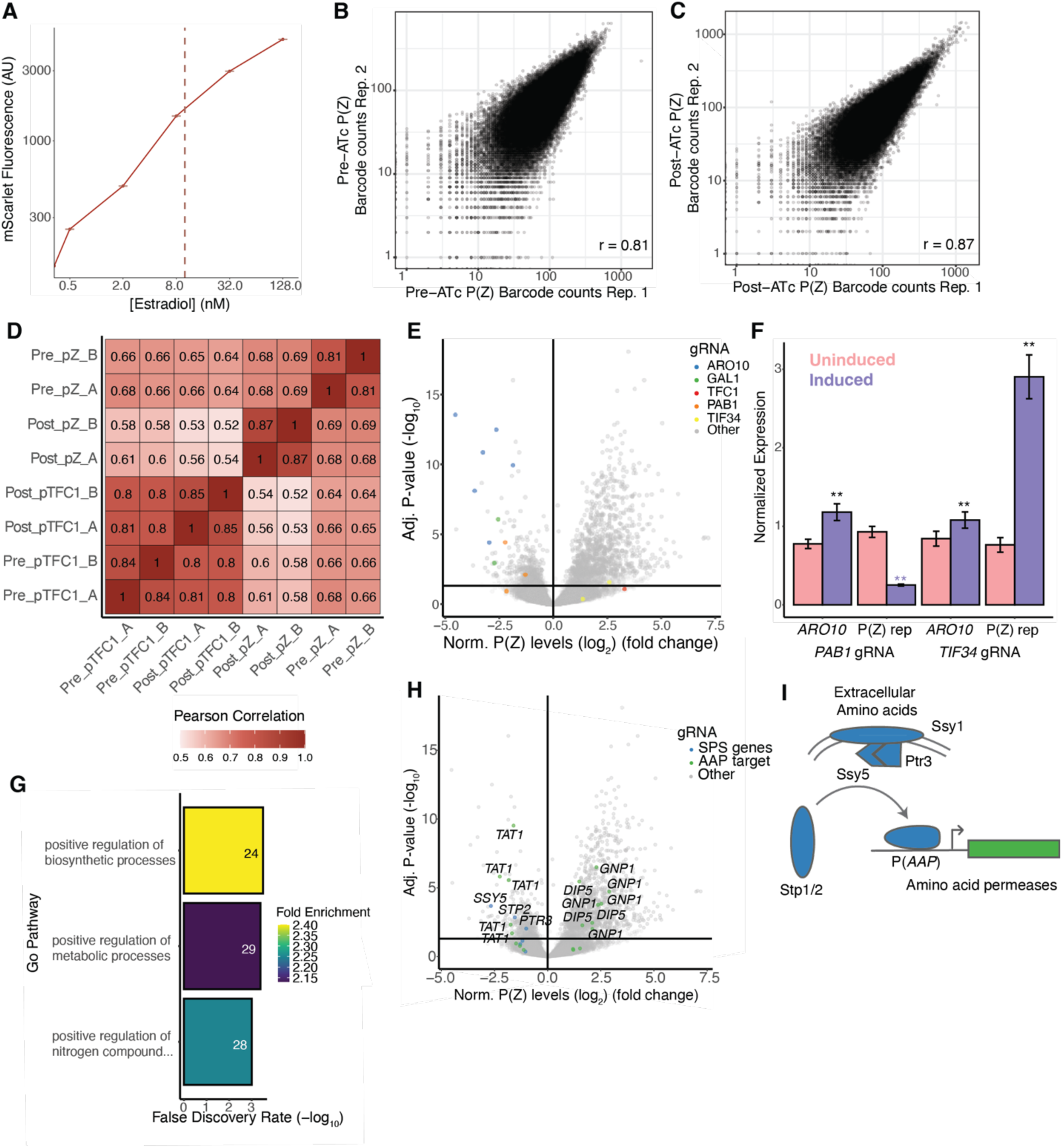
CiBER-seq profile *ARO10* expression regulation. **(A)**Titration across a range of β-estradiol concentrations in *ARO10-P2A-ZEM* cells to determine a concentration in the linear range where mScarlet fluorescence. Median fluorescence values and standard error are plotted. Line indicates the concentration chosen for CiBER-seq screen. **(B)** Comparison of replicate P(Z) barcode codes from pre-gRNA induction samples and corresponding correlation coefficient (Pearson’s). These were chosen as a representative example of replicate samples. **(C)** Same as in (B) for post-gRNA induction samples. **(D)** Pearson’s correlation coefficients of pairwise comparisons across all RNA-sequencing samples. Color represents magnitude of correlation coefficient. **(E)** CiBER-seq results showing fold-change (log2) in P(Z) reporter abundance, relative to P(*TFC1*) reporter levels, for each gRNA. Line indicates significance cutoff (adjusted p-value < 0.05). Color represents gene identity. **(F)** RT-qPCR of *ARO10* and mScarlet reporter expression following *PAB1* or *TIF34* gRNA induction. (**) represents p < 0.05 calculated by Student’s t test. **(G)** GO analysis for genes targeted by gRNAs that down-regulated P(Z) expression. gRNAs were filtered for fold-change (log2) > 1 and adjusted p-value < 0.05. The most statistically significant entries were chosen and narrowed based on percentage of overlapping genes. **(H)** Same as in (E) except highlight components of the SPS amino acid sensing pathway and target amino acid permease genes. **(I)** Schematic of the SPS amino acid sensing pathway.

**Figure S5:**
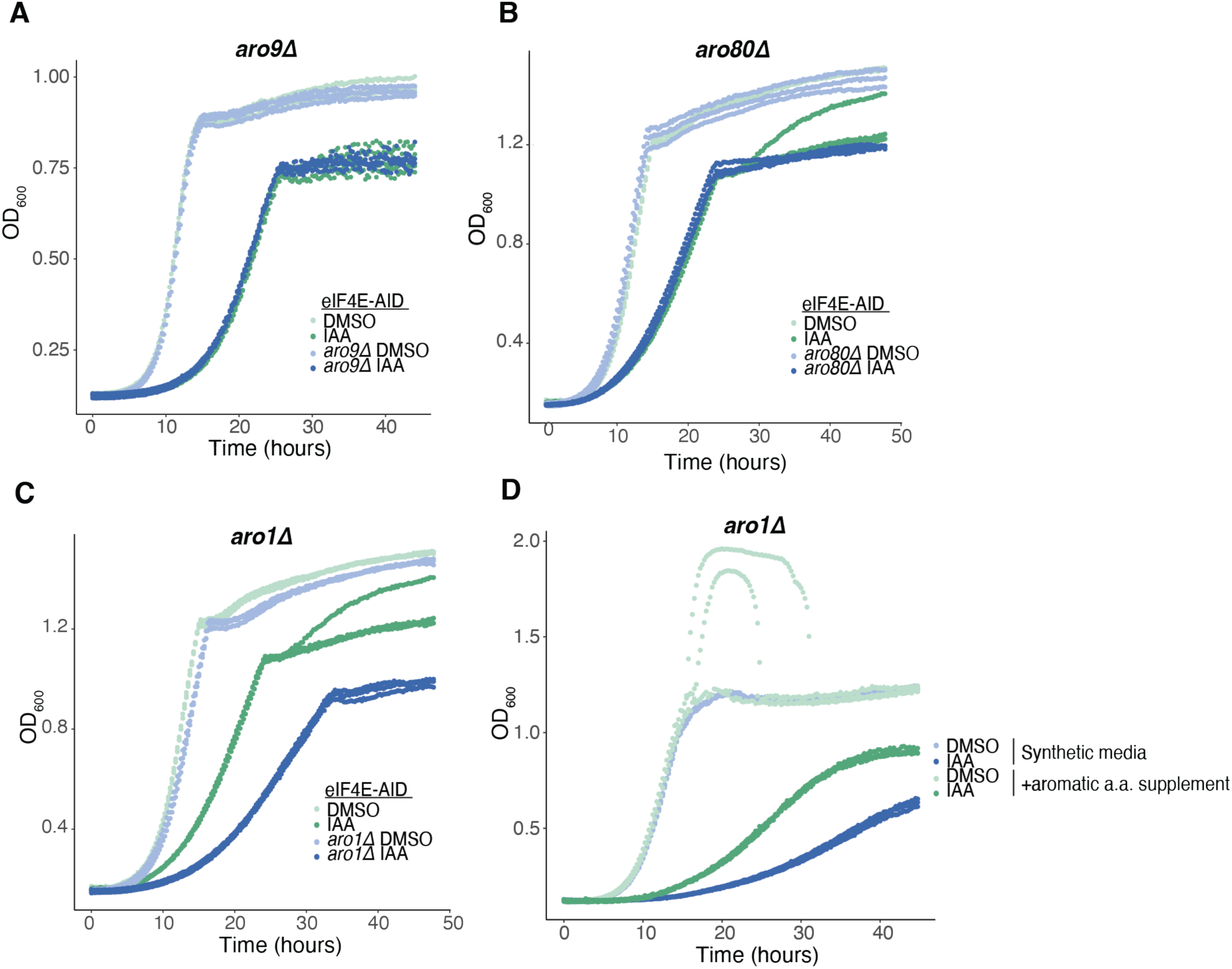
Futile cycles of aromatic amino acid metabolism. **(A)** Growth curves of eIF4E-AID and eIF4E-AID *aro9Δ* cells maintain in IAA or DMSO. **(B)** Same as in (A) except for *aro80Δ* cells. **(C)** Same as in (A) except for *aro1Δ* cells. **(D)** Growth curves of eIF4E-AID *aro1Δ* cells maintain in IAA or DMSO in synthetic complete media with and without aromatic amino acid supplementation.

**Figure S6:**
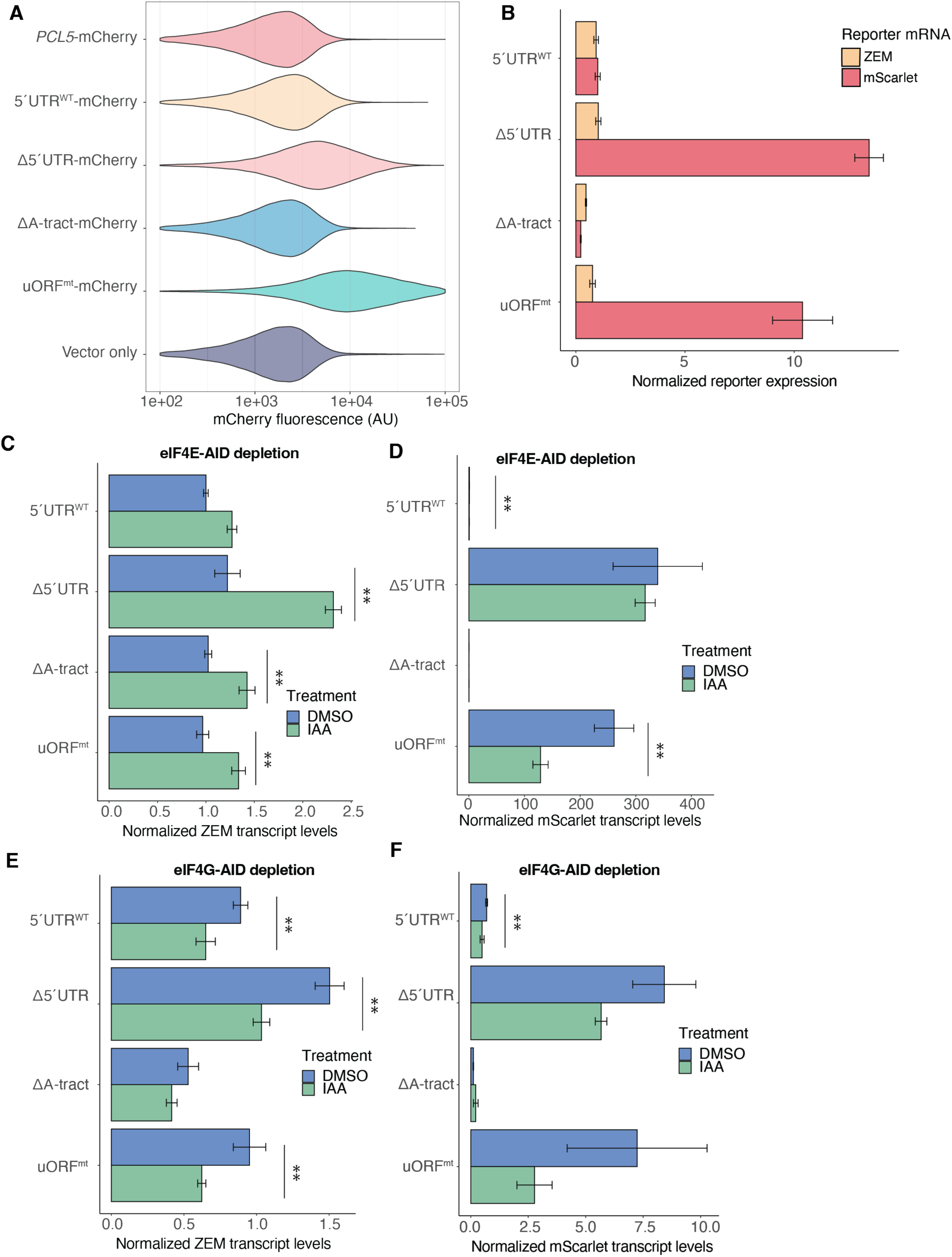
Translation regulation of *PCL5.* **(A)** Distribution of fluorescence of *PCL5*-mCherry reporters and vector only control. **(B)** RT-qPCR of ZEM transcript and mScarlet reporter expression of *PCL5* 5’UTR. Error bars represent standard deviation, n=3. **(C)** RT-qPCR of ZEM transcript expression of *PCL5* 5’UTR mutants following eIF4E-AID depletion. (**) represents p < 0.05 calculated by Student’s t test. Error bars represent standard deviation, n=3. **(D)** Same as in (C) except mScarlet reporter transcript. **(E)** Same as in (C) except in eIF4G-AID depletion. **(F)** Same as in (E) except mScarlet reporter transcript.

**Figure S7:**
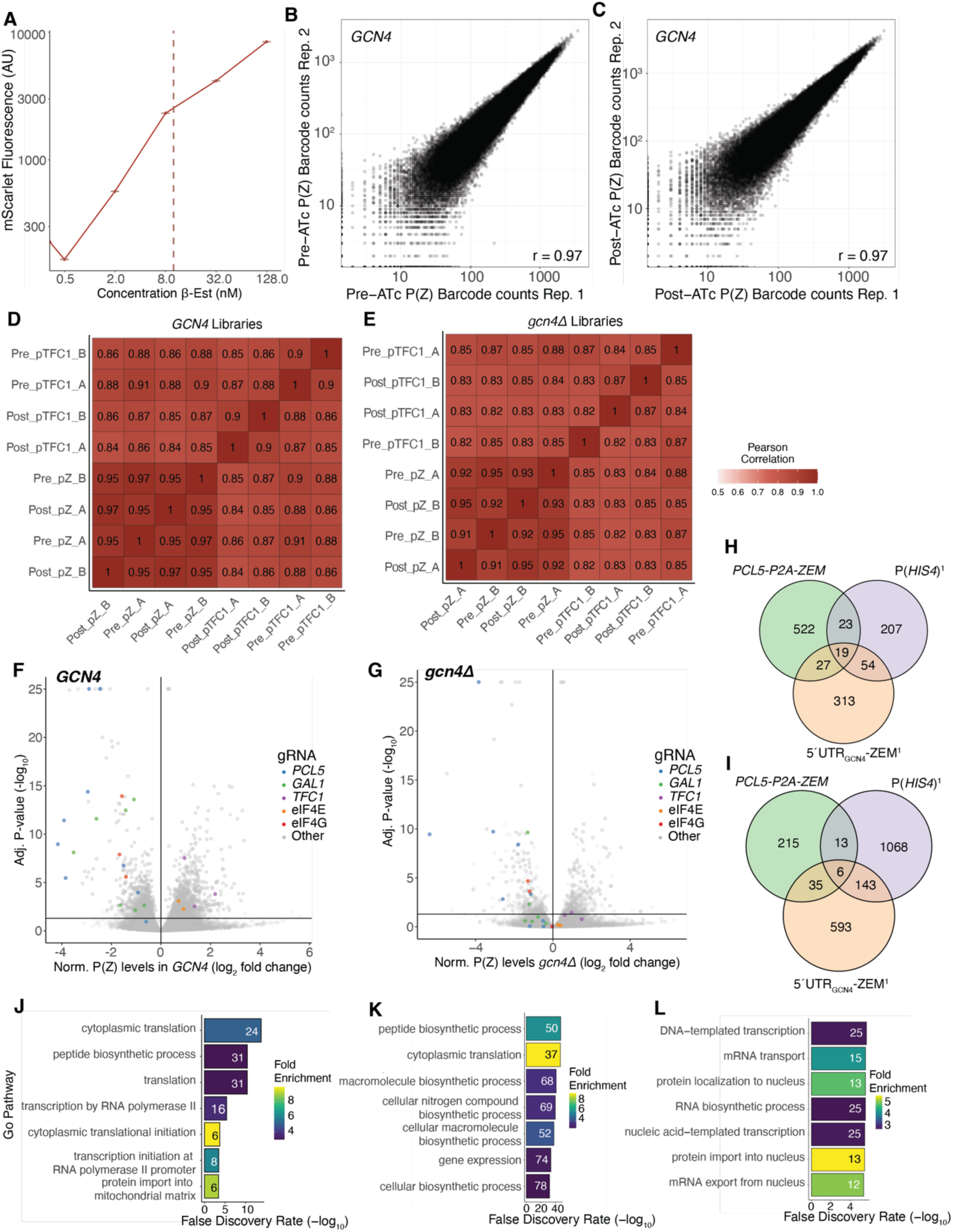
CiBER-seq profile *PCL5* expression regulation. **(A)** Titration across a range of β-estradiol concentrations in *PCL5-P2A-ZEM* cells to determine a concentration in the linear range where mScarlet fluorescence. Median fluorescence values and standard error are plotted. Line indicates the concentration chosen for CiBER-seq screen. **(B)** Comparison of replicate P(Z) barcode codes from pre-gRNA induction samples and corresponding correlation coefficient (Pearson’s). These were chosen as a representative example of replicate samples. **(C)** Same as in (B) for post-gRNA induction samples. **(D)** Pearson’s correlation coefficients of pairwise comparisons across all *GCN4* samples. Color represents magnitude of correlation coefficient. **(E)** Same as in (D) except for *gcn4Δ* samples. **(F)** CiBER-seq results showing fold-change (log2) in P(Z) reporter abundance in *GCN4* cells, relative to P(*TFC1*) reporter levels, for each gRNA. Line indicates significance cutoff (adjusted p-value < 0.05). Color represents gene identity. **(G)** Same as in (F) except in *gcn4Δ* cells. **(H)** Overlap between gRNAs that produced significant (adjusted p-value < 0.05) and substantial increase (fold-change (log2) > 1) in *PCL5*-P2A-ZEM (this study), P(*HIS4),* and 5’UTRGCN4-ZEM reporters expression in previously published CiBER-seq analysis of *GCN4* (Muller et al., 2020). **(I)** Same as in (H) except for gRNAs that produced a substantial decrease in reporter expression. **(J)** GO analysis for genes targeted by gRNAs that up-regulated P(Z) expression in *GCN4* cells. gRNAs were filtered for fold-change (log2) > 1 and adjusted p-value < 0.05. The most statistically significant entries were chosen and narrowed based on percentage of overlapping genes. **(K)** Same as in (J) for *gcn4Δ* cells. **(L)** Same as in (K) for gRNAs that down-regulated P(Z) expression.

**Figure S8:**
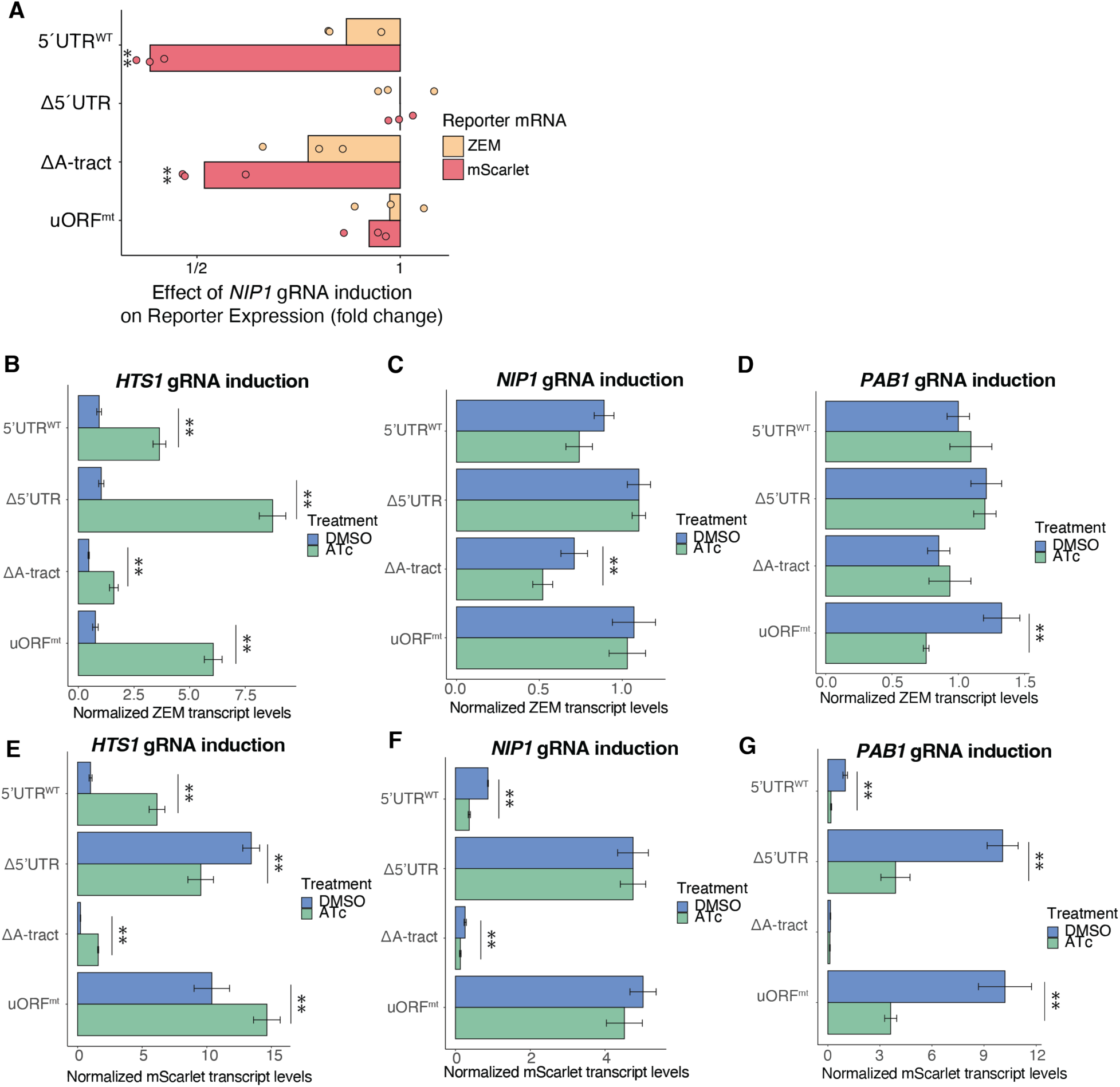
Trans factor regulation of *PCL5* translation. **(A)** RT-qPCR of ZEM transcript and mScarlet reporter expression of *PCL5* 5’UTR mutants following HTS1 gRNA induction. Individual reporters were normalized to uninduced control. (**) represents p < 0.05 calculated by Student’s t test. **(B)** RT-qPCR of ZEM transcript expression of *PCL5* 5’UTR mutants following induction of *HTS1* gRNA. (**) represents p < 0.05 calculated by Student’s t test. Error bars represent standard deviation, n=3. **(C)** Same as in (B) except following *NIP1* gRNA induction. **(D)** Same as in (B) except following *PAB1* gRNA induction. **(E)** RT-qPCR of mScarlet transcript expression of *PCL5* 5’UTR mutants following induction of *HTS1* gRNA. **(F)** Same as in (E) except following *NIP1* gRNA induction. **(G)** Same as in (E) except following *PAB1* gRNA induction.

## References

Abraham, J., Bhat, S.G., 2008. Permeabilization of baker’s yeast with N-lauroyl sarcosine. J Ind Microbiol Biotechnol 35, 799–804. https://doi.org/10.1007/s10295-008-0350-9

Amrani, N., Ganesan, R., Kervestin, S., Mangus, D.A., Ghosh, S., Jacobson, A., 2004. A faux 3′-UTR promotes aberrant termination and triggers nonsense-mediated mRNA decay. Nature 432, 112–118. https://doi.org/10.1038/nature03060

Amrani, N., Ghosh, S., Mangus, D.A., Jacobson, A., 2008. Translation factors promote the formation of two states of the closed-loop mRNP. Nature 453, 1276–1280. https://doi.org/10.1038/nature06974

Aranda-Díaz, A., Mace, K., Zuleta, I., Harrigan, P., El-Samad, H., 2017. Robust Synthetic Circuits for Two-Dimensional Control of Gene Expression in Yeast. ACS Synth. Biol. 6, 545–554. https://doi.org/10.1021/acssynbio.6b00251

Ares, M., 2012. Isolation of Total RNA from Yeast Cell Cultures: Figure 1. Cold Spring Harb Protoc 2012, pdb.prot071456. https://doi.org/10.1101/pdb.prot071456

Ashe, M.P., 2001. A novel eIF2B-dependent mechanism of translational control in yeast as a response to fusel alcohols. The EMBO Journal 20, 6464–6474. https://doi.org/10.1093/emboj/20.22.6464

Ashuach, T., Fischer, D.S., Kreimer, A., Ahituv, N., Theis, F.J., Yosef, N., 2019. MPRAnalyze: statistical framework for massively parallel reporter assays. Genome Biol 20, 183. https://doi.org/10.1186/s13059-019-1787-z

Bajmoczi, M., Sneve, M., Eide, D.J., Drewes, L.R., 1998. TAT1 Encodes a Low-Affinity Histidine Transporter inSaccharomyces cerevisiae. Biochemical and Biophysical Research Communications 243, 205–209. https://doi.org/10.1006/bbrc.1998.8082

Brar, G.A., Yassour, M., Friedman, N., Regev, A., Ingolia, N.T., Weissman, J.S., 2012. High-Resolution View of the Yeast Meiotic Program Revealed by Ribosome Profiling. Science 335, 552–557. https://doi.org/10.1126/science.1215110

Brothers, M., Rine, J., 2019. Mutations in the PCNA DNA Polymerase Clamp of *Saccharomyces cerevisiae* Reveal Complexities of the Cell Cycle and Ploidy on Heterochromatin Assembly. Genetics 213, 449–463. https://doi.org/10.1534/genetics.119.302452

Calkhoven, C.F., Müller, C., Leutz, A., 2000. Translational control of C/EBPα and C/EBPβ isoform expression. Genes Dev. 14, 1920–1932. https://doi.org/10.1101/gad.14.15.1920

Çetin, B., O’Leary, S.E., 2022. mRNA- and factor-driven dynamic variability controls eIF4F-cap recognition for translation initiation. Nucleic Acids Research 50, 8240– 8261. https://doi.org/10.1093/nar/gkac631

Chan, L.Y., Mugler, C.F., Heinrich, S., Vallotton, P., Weis, K., 2018. Non-invasive measurement of mRNA decay reveals translation initiation as the major determinant of mRNA stability. eLife 7, e32536. https://doi.org/10.7554/eLife.32536

Chen, H., Fink, G.R., 2006. Feedback control of morphogenesis in fungi by aromatic alcohols. Genes Dev. 20, 1150–1161. https://doi.org/10.1101/gad.1411806

Costello, J., Castelli, L.M., Rowe, W., Kershaw, C.J., Talavera, D., Mohammad-Qureshi, S.S., Sims, P.F.G., Grant, C.M., Pavitt, G.D., Hubbard, S.J., Ashe, M.P., 2015. Global mRNA selection mechanisms for translation initiation. Genome Biol 16, 10. https://doi.org/10.1186/s13059-014-0559-z

Cox, J.S., Chapman, R.E., Walter, P., 1997. The unfolded protein response coordinates the production of endoplasmic reticulum protein and endoplasmic reticulum membrane. MBoC 8, 1805–1814. https://doi.org/10.1091/mbc.8.9.1805

Crawford, R.A., Pavitt, G.D., 2019. Translational regulation in response to stress in *Saccharomyces cerevisiae*. Yeast 36, 5–21. https://doi.org/10.1002/yea.3349

Cvijović, M., Dalevi, D., Bilsland, E., Kemp, G.J., Sunnerhagen, P., 2007. Identification of putative regulatory upstream ORFs in the yeast genome using heuristics and evolutionary conservation. BMC Bioinformatics 8, 295. https://doi.org/10.1186/1471-2105-8-295

Danaie, P., Altmann, M., Hall, M.N., Trachsel, H., Helliwell, S.B., 1999. CLN3 expression is sufficient to restore G1-to-S-phase progression in Saccharomyces cerevisiae mutants defective in translation initiation factor eIF4E. Biochemical Journal 340, 135–141. https://doi.org/10.1042/bj3400135

Dever, T.E., Feng, L., Wek, R.C., Cigan, A.M., Donahue, T.F., Hinnebusch, A.G., 1992. Phosphorylation of initiation factor 2α by protein kinase GCN2 mediates gene-specific translational control of GCN4 in yeast. Cell 68, 585–596. https://doi.org/10.1016/0092-8674(92)90193-G

Dobin, A., Davis, C.A., Schlesinger, F., Drenkow, J., Zaleski, C., Jha, S., Batut, P., Chaisson, M., Gingeras, T.R., 2013. STAR: ultrafast universal RNA-seq aligner. Bioinformatics 29, 15–21. https://doi.org/10.1093/bioinformatics/bts635

Duncan, K., Edwards, R.M., Coggins, J.R., 1988. The Saccharomyces cerevisiae ARO1 gene. An example of the co-ordinate regulation of five enzymes on a single biosynthetic pathway. FEBS Lett 241, 83–88. https://doi.org/10.1016/0014-5793(88)81036-6

Eckert-Boulet, N., Nielsen, P.S., Friis, C., dos Santos, M.M., Nielsen, J., Kielland-Brandt, M.C., Regenberg, B., 2004. Transcriptional profiling of extracellular amino acid sensing in *Saccharomyces cerevisiae* and the role of Stp1p and Stp2p. Yeast 21, 635–648. https://doi.org/10.1002/yea.1120

Firczuk, H., Kannambath, S., Pahle, J., Claydon, A., Beynon, R., Duncan, J., Westerhoff, H., Mendes, P., McCarthy, J.E., 2013. An *in vivo* control map for the eukaryotic mRNA translation machinery. Mol Syst Biol 9, 635. https://doi.org/10.1038/msb.2012.73

Fonseca, B.D., Zakaria, C., Jia, J.-J., Graber, T.E., Svitkin, Y., Tahmasebi, S., Healy, D., Hoang, H.-D., Jensen, J.M., Diao, I.T., Lussier, A., Dajadian, C., Padmanabhan, N., Wang, W., Matta-Camacho, E., Hearnden, J., Smith, E.M., Tsukumo, Y., Yanagiya, A., Morita, M., Petroulakis, E., González, J.L., Hernández, G., Alain, T., Damgaard, C.K., 2015. La-related Protein 1 (LARP1) Represses Terminal Oligopyrimidine (TOP) mRNA Translation Downstream of mTOR Complex 1 (mTORC1). Journal of Biological Chemistry 290, 15996–16020. https://doi.org/10.1074/jbc.M114.621730

Gibson, D.G., Young, L., Chuang, R.-Y., Venter, J.C., Hutchison, C.A., Smith, H.O., 2009. Enzymatic assembly of DNA molecules up to several hundred kilobases. Nat Methods 6, 343–345. https://doi.org/10.1038/nmeth.1318

Gietz, R.D., Schiestl, R.H., 2007. High-efficiency yeast transformation using the LiAc/SS carrier DNA/PEG method. Nat Protoc 2, 31–34. https://doi.org/10.1038/nprot.2007.13

Gilbert, W.V., Zhou, K., Butler, T.K., Doudna, J.A., 2007. Cap-Independent Translation Is Required for Starvation-Induced Differentiation in Yeast. Science 317, 1224– 1227. https://doi.org/10.1126/science.1144467

Goldstein, A.L., McCusker, J.H., 1999. Three new dominant drug resistance cassettes for gene disruption inSaccharomyces cerevisiae. Yeast 15, 1541–1553. https://doi.org/10.1002/(SICI)1097-0061(199910)15:14<1541::AID-YEA476>3.0.CO;2-K

Groušl, T., Ivanov, P., Frydlová, I., Vašicová, P., Janda, F., Vojtová, J., Malínská, K., Malcová, I., Nováková, L., Janošková, D., Valášek, L., Hašek, J., 2009. Robust heat shock induces eIF2α-phosphorylation-independent assembly of stress granules containing eIF3 and 40S ribosomal subunits in budding yeast, *Saccharomyces cerevisiae*. Journal of Cell Science 122, 2078–2088. https://doi.org/10.1242/jcs.045104

Hazelwood, L.A., Daran, J.-M., van Maris, A.J.A., Pronk, J.T., Dickinson, J.R., 2008. The Ehrlich Pathway for Fusel Alcohol Production: a Century of Research on *Saccharomyces cerevisiae* Metabolism. Appl Environ Microbiol 74, 3920–3920. https://doi.org/10.1128/AEM.00934-08

Hentges, P., Van Driessche, B., Tafforeau, L., Vandenhaute, J., Carr, A.M., 2005. Three novel antibiotic marker cassettes for gene disruption and marker switching inSchizosaccharomyces pombe. Yeast 22, 1013–1019. https://doi.org/10.1002/yea.1291

Hershey, J.W.B., Sonenberg, N., Mathews, M.B., 2012. Principles of Translational Control: An Overview. Cold Spring Harbor Perspectives in Biology 4, a011528– a011528. https://doi.org/10.1101/cshperspect.a011528

Hinnebusch, A.G., 2005. TRANSLATIONAL REGULATION OF *GCN4* AND THE GENERAL AMINO ACID CONTROL OF YEAST. Annu. Rev. Microbiol. 59, 407–450. https://doi.org/10.1146/annurev.micro.59.031805.133833

Ingolia, N.T., Brar, G.A., Rouskin, S., McGeachy, A.M., Weissman, J.S., 2012. The ribosome profiling strategy for monitoring translation in vivo by deep sequencing of ribosome-protected mRNA fragments. Nat Protoc 7, 1534–1550. https://doi.org/10.1038/nprot.2012.086

Ingolia, N.T., Ghaemmaghami, S., Newman, J.R.S., Weissman, J.S., 2009. Genome-Wide Analysis in Vivo of Translation with Nucleotide Resolution Using Ribosome Profiling. Science 324, 218–223. https://doi.org/10.1126/science.1168978

Krause, L., Willing, F., Andreou, A.Z., Klostermeier, D., 2022. The domains of yeast eIF4G, eIF4E and the cap fine-tune eIF4A activities through an intricate network of stimulatory and inhibitory effects. Nucleic Acids Research 50, 6497–6510. https://doi.org/10.1093/nar/gkac437

Kubota, T., Nishimura, K., Kanemaki, M.T., Donaldson, A.D., 2013. The Elg1 Replication Factor C-like Complex Functions in PCNA Unloading during DNA Replication. Molecular Cell 50, 273–280. https://doi.org/10.1016/j.molcel.2013.02.012

Lahr, R.M., Fonseca, B.D., Ciotti, G.E., Al-Ashtal, H.A., Jia, J.-J., Niklaus, M.R., Blagden, S.P., Alain, T., Berman, A.J., 2017. La-related protein 1 (LARP1) binds the mRNA cap, blocking eIF4F assembly on TOP mRNAs. eLife 6, e24146. https://doi.org/10.7554/eLife.24146

Lee, K., Hahn, J.-S., 2013. Interplay of Aro80 and GATA activators in regulation of genes for catabolism of aromatic amino acids in *Saccharomyces cerevisiae*: Regulation of genes for aromatic amino acids catabolism. Molecular Microbiology 88, 1120–1134. https://doi.org/10.1111/mmi.12246

Lee, M.E., DeLoache, W.C., Cervantes, B., Dueber, J.E., 2015. A Highly Characterized Yeast Toolkit for Modular, Multipart Assembly. ACS Synth. Biol. 4, 975–986. https://doi.org/10.1021/sb500366v

Liu, B., Qian, S.-B., 2014. Translational reprogramming in cellular stress response: Translational reprogramming in stress. WIREs RNA 5, 301–305. https://doi.org/10.1002/wrna.1212

Ljungdahl, P.O., 2009. Amino-acid-induced signalling via the SPS-sensing pathway in yeast. Biochemical Society Transactions 37, 242–247. https://doi.org/10.1042/BST0370242

Love, M.I., Huber, W., Anders, S., 2014. Moderated estimation of fold change and dispersion for RNA-seq data with DESeq2. Genome Biol 15, 550. https://doi.org/10.1186/s13059-014-0550-8

Lu, P.D., Harding, H.P., Ron, D., 2004. Translation reinitiation at alternative open reading frames regulates gene expression in an integrated stress response. Journal of Cell Biology 167, 27–33. https://doi.org/10.1083/jcb.200408003

Marcotrigiano, J., Gingras, A.-C., Sonenberg, N., Burley, S.K., 1997. Cocrystal Structure of the Messenger RNA 5′ Cap-Binding Protein (eIF4E) Bound to 7-methyl-GDP. Cell 89, 951–961. https://doi.org/10.1016/S0092-8674(00)80280-9

Martin, M., 2011. Cutadapt removes adapter sequences from high-throughput sequencing reads. EMBnet j. 17, 10. https://doi.org/10.14806/ej.17.1.200

Matsuo, H., Li, H., McGuire, A.M., Fletcher, C.M., Gingras, A.-C., Sonenberg, N., Wagner, G., 1997. Structure of translation factor elF4E bound to m7GDP and interaction with 4E-binding protein. Nat Struct Mol Biol 4, 717–724. https://doi.org/10.1038/nsb0997-717

McGeachy, A.M., Meacham, Z.A., Ingolia, N.T., 2019. An Accessible Continuous-Culture Turbidostat for Pooled Analysis of Complex Libraries. ACS Synth. Biol. 8, 844–856. https://doi.org/10.1021/acssynbio.8b00529

McGlincy, N.J., Ingolia, N.T., 2017. Transcriptome-wide measurement of translation by ribosome profiling. Methods 126, 112–129. https://doi.org/10.1016/j.ymeth.2017.05.028

Meyuhas, O., Kahan, T., 2015. The race to decipher the top secrets of TOP mRNAs. Biochimica et Biophysica Acta (BBA) - Gene Regulatory Mechanisms 1849, 801–811. https://doi.org/10.1016/j.bbagrm.2014.08.015

Mueller, P.P., Hinnebusch, A.G., 1986. Multiple upstream AUG codons mediate translational control of GCN4. Cell 45, 201–207. https://doi.org/10.1016/0092-8674(86)90384-3

Muller, R., Meacham, Z., Ingolia, N., 2022. Plasmid and Sequencing Library Preparation for CRISPRi Barcoded Expression Reporter Sequencing (CiBER-seq) in Saccharomyces cerevisiae. BIO-PROTOCOL 12. https://doi.org/10.21769/BioProtoc.4376

Muller, R., Meacham, Z.A., Ferguson, L., Ingolia, N.T., 2020. CiBER-seq dissects genetic networks by quantitative CRISPRi profiling of expression phenotypes. Science 370, eabb9662. https://doi.org/10.1126/science.abb9662

Nishimura, K., Fukagawa, T., Takisawa, H., Kakimoto, T., Kanemaki, M., 2009. An auxin-based degron system for the rapid depletion of proteins in nonplant cells. Nat Methods 6, 917–922. https://doi.org/10.1038/nmeth.1401

O’Leary, S.E., Petrov, A., Chen, J., Puglisi, J.D., 2013. Dynamic Recognition of the mRNA Cap by Saccharomyces cerevisiae eIF4E. Structure 21, 2197–2207. https://doi.org/10.1016/j.str.2013.09.016

Park, E.-H., Zhang, F., Warringer, J., Sunnerhagen, P., Hinnebusch, A.G., 2011. Depletion of eIF4G from yeast cells narrows the range of translational efficiencies genome-wide. BMC Genomics 12, 68. https://doi.org/10.1186/1471-2164-12-68

Philippe, L., van den Elzen, A.M.G., Watson, M.J., Thoreen, C.C., 2020. Global analysis of LARP1 translation targets reveals tunable and dynamic features of 5′ TOP motifs. Proc. Natl. Acad. Sci. U.S.A. 117, 5319–5328. https://doi.org/10.1073/pnas.1912864117

Reid, B.J., Hartwell, L.H., 1977. Regulation of mating in the cell cycle of Saccharomyces cerevisiae. Journal of Cell Biology 75, 355–365. https://doi.org/10.1083/jcb.75.2.355

Riback, J.A., Katanski, C.D., Kear-Scott, J.L., Pilipenko, E.V., Rojek, A.E., Sosnick, T.R., Drummond, D.A., 2017. Stress-Triggered Phase Separation Is an Adaptive, Evolutionarily Tuned Response. Cell 168, 1028–1040.e19. https://doi.org/10.1016/j.cell.2017.02.027

Sachs, A.B., Davis, R.W., Kornberg, R.D., 1987. A single domain of yeast poly(A)-binding protein is necessary and sufficient for RNA binding and cell viability. Mol Cell Biol 7, 3268–3276. https://doi.org/10.1128/mcb.7.9.3268-3276.1987

Schmidt, A., Hall, M.N., Koller, A., 1994. Two FK506 resistance-conferring genes in Saccharomyces cerevisiae, TAT1 and TAT2, encode amino acid permeases mediating tyrosine and tryptophan uptake. Mol Cell Biol 14, 6597–6606. https://doi.org/10.1128/mcb.14.10.6597-6606.1994

Sen, N.D., Zhou, F., Harris, M.S., Ingolia, N.T., Hinnebusch, A.G., 2016. eIF4B stimulates translation of long mRNAs with structured 5′ UTRs and low closed-loop potential but weak dependence on eIF4G. Proc. Natl. Acad. Sci. U.S.A. 113, 10464–10472. https://doi.org/10.1073/pnas.1612398113

Shah, P., Ding, Y., Niemczyk, M., Kudla, G., Plotkin, J.B., 2013. Rate-Limiting Steps in Yeast Protein Translation. Cell 153, 1589–1601. https://doi.org/10.1016/j.cell.2013.05.049

Shemer, R., Meimoun, A., Holtzman, T., Kornitzer, D., 2002. Regulation of the Transcription Factor Gcn4 by Pho85 Cyclin Pcl5. Molecular and Cellular Biology 22, 5395–5404. https://doi.org/10.1128/MCB.22.15.5395-5404.2002

Sonenberg, N., Hinnebusch, A.G., 2009. Regulation of Translation Initiation in Eukaryotes: Mechanisms and Biological Targets. Cell 136, 731–745. https://doi.org/10.1016/j.cell.2009.01.042

Sprouffske, K., Wagner, A., 2016. Growthcurver: an R package for obtaining interpretable metrics from microbial growth curves. BMC Bioinformatics 17, 172. https://doi.org/10.1186/s12859-016-1016-7

Staschke, K.A., Dey, S., Zaborske, J.M., Palam, L.R., McClintick, J.N., Pan, T., Edenberg, H.J., Wek, R.C., 2010. Integration of General Amino Acid Control and Target of Rapamycin (TOR) Regulatory Pathways in Nitrogen Assimilation in Yeast. Journal of Biological Chemistry 285, 16893–16911. https://doi.org/10.1074/jbc.M110.121947

Steffen, K.K., MacKay, V.L., Kerr, E.O., Tsuchiya, M., Hu, D., Fox, L.A., Dang, N., Johnston, E.D., Oakes, J.A., Tchao, B.N., Pak, D.N., Fields, S., Kennedy, B.K., Kaeberlein, M., 2008. Yeast Life Span Extension by Depletion of 60S Ribosomal Subunits Is Mediated by Gcn4. Cell 133, 292–302. https://doi.org/10.1016/j.cell.2008.02.037

Stovicek, V., Borja, G.M., Forster, J., Borodina, I., 2015. EasyClone 2.0: expanded toolkit of integrative vectors for stable gene expression in industrial *Saccharomyces cerevisiae* strains. Journal of Industrial Microbiology and Biotechnology 42, 1519–1531. https://doi.org/10.1007/s10295-015-1684-8

Thompson, M.K., Gilbert, W.V., 2017. mRNA length-sensing in eukaryotic translation: reconsidering the “closed loop” and its implications for translational control. Curr Genet 63, 613–620. https://doi.org/10.1007/s00294-016-0674-3

Thoreen, C.C., Chantranupong, L., Keys, H.R., Wang, T., Gray, N.S., Sabatini, D.M., 2012. A unifying model for mTORC1-mediated regulation of mRNA translation. Nature 485, 109–113. https://doi.org/10.1038/nature11083

Truitt, M.L., Conn, C.S., Shi, Z., Pang, X., Tokuyasu, T., Coady, A.M., Seo, Y., Barna, M., Ruggero, D., 2015. Differential Requirements for eIF4E Dose in Normal Development and Cancer. Cell 162, 59–71. https://doi.org/10.1016/j.cell.2015.05.049

Vilela, C., 2000. The eukaryotic mRNA decapping protein Dcp1 interacts physically and functionally with the eIF4F translation initiation complex. The EMBO Journal 19, 4372–4382. https://doi.org/10.1093/emboj/19.16.4372

von der Haar, T., Gross, J.D., Wagner, G., McCarthy, J.E.G., 2004. The mRNA cap-binding protein eIF4E in post-transcriptional gene expression. Nat Struct Mol Biol 11, 503–511. https://doi.org/10.1038/nsmb779

von der Haar, T., McCarthy, J.E.G., 2002. Intracellular translation initiation factor levels in *Saccharomyces cerevisiae* and their role in cap-complex function: Translation initiation factor levels in yeast. Molecular Microbiology 46, 531–544. https://doi.org/10.1046/j.1365-2958.2002.03172.x

Vopálenský, V., Sýkora, M., Mašek, T., Pospíšek, M., 2019. Messenger RNAs of Yeast Virus-Like Elements Contain Non-templated 5′ Poly(A) Leaders, and Their Expression Is Independent of eIF4E and Pab1. Front. Microbiol. 10, 2366. https://doi.org/10.3389/fmicb.2019.02366

Wiltschi, B., Wenger, W., Nehring, S., Budisa, N., 2008. Expanding the genetic code of *Saccharomyces cerevisiae* with methionine analogues. Yeast 25, 775–786. https://doi.org/10.1002/yea.1632

Wu, C.C.-C., Peterson, A., Zinshteyn, B., Regot, S., Green, R., 2020. Ribosome Collisions Trigger General Stress Responses to Regulate Cell Fate. Cell 182, 404–416.e14. https://doi.org/10.1016/j.cell.2020.06.006

Xia, H., Shangguan, L., Chen, S., Yang, Q., Zhang, X., Yao, L., Yang, S., Dai, J., Chen, X., 2022. Rapamycin enhanced the production of 2-phenylethanol during whole-cell bioconversion by yeast. Appl Microbiol Biotechnol 106, 6471–6481. https://doi.org/10.1007/s00253-022-12169-6

Young, S.K., Wek, R.C., 2016. Upstream Open Reading Frames Differentially Regulate Gene-specific Translation in the Integrated Stress Response. Journal of Biological Chemistry 291, 16927–16935. https://doi.org/10.1074/jbc.R116.733899

Zhang, D., Wang, F., Yu, Y., Ding, S., Chen, T., Sun, W., Liang, C., Yu, B., Ying, H., Liu, D., Chen, Y., 2021. Effect of quorum-sensing molecule 2-phenylethanol and ARO genes on Saccharomyces cerevisiae biofilm. Appl Microbiol Biotechnol 105, 3635–3648. https://doi.org/10.1007/s00253-021-11280-4

Zhang, Z., 2005. Mapping of transcription start sites in Saccharomyces cerevisiae using 5’ SAGE. Nucleic Acids Research 33, 2838–2851. https://doi.org/10.1093/nar/gki583

Zinshteyn, B., Rojas-Duran, M.F., Gilbert, W.V., 2017. Translation initiation factor eIF4G1 preferentially binds yeast transcript leaders containing conserved oligo-uridine motifs. RNA 23, 1365–1375. https://doi.org/10.1261/rna.062059.117

Zou, K., Rouskin, S., Dervishi, K., McCormick, M.A., Sasikumar, A., Deng, C., Chen, Z., Kaeberlein, M., Brem, R.B., Polymenis, M., Kennedy, B.K., Weissman, J.S., Zheng, J., Ouyang, Q., Li, H., 2020. Life span extension by glucose restriction is abrogated by methionine supplementation: Cross-talk between glucose and methionine and implication of methionine as a key regulator of life span. Sci. Adv. 6, eaba1306. https://doi.org/10.1126/sciadv.aba1306

